# Predicting the tension in actin cytoskeleton from the nucleus shape

**DOI:** 10.1101/2020.08.28.272435

**Authors:** Sreenath Balakrishnan, Shilpa R Raju, Anwesha Barua, G.K. Ananthasuresh

## Abstract

Tension in actin cytoskeleton regulates many cellular processes and nuclear morphology. Here, we demonstrate a simple computational method for estimating actin cytoskeletal tension from nucleus shape. We first note that mechanics-based modeling defines a relationship among the volume, surface area, and projected area of the nucleus and hence a specific surface in the three-parameter space of the aforementioned geometric quantities. Data of nuclei from multiple cell types lie on such a surface. Furthermore, nuclei from a given cell population lie on a straight line on the surface. The location and orientation of the line varies with cell type. By using a mechanical model, we present two non-dimensional parameters, namely, the flatness and stretch indicators, which serve as curvilinear coordinates on the surface. Flatness indicator defines the extent of nuclear flattening due to actin cytoskeletal tension and the stretch indicator captures the effect of the elastic modulus of the nuclear envelope. We validate our assertions by modulating the actin cytoskeletal tension using three independent mechanisms: (i) direct downregulation by Cytochalasin D, (ii) indirect upregulation using Nocodazole, and (iii) mechanical stimulation by varying substrate stiffness. We also infer that the flatness indicator is equivalent to the ratio of the height to diameter of the nucleus and is related to the Vogel number. By using this geometric insight, we validate the predictions of our model with data from many previous studies. Finally, we present an analytical formula and a correlation for estimating actin cytoskeletal tension from nuclear projected area and volume.

## Introduction

Nucleus morphology is an indicator of cellular function and dysfunction. Many diseases such as cancer (1, 2) and laminopathies (3–5) are known to alter nucleus shape. Aberrations in nuclear shape were directly correlated with these diseases and used as a marker for disease diagnosis. It is known that diseases modify nuclear shape indirectly by modulating one or more cellular factors governing nuclear morphology such as cytoskeleton (6), lamins (7), chromatin (8), and osmotic pressure difference across the nuclear envelope (9, 10). Hence for broad-range disease diagnosis with specificity, it is beneficial to have a technique for predicting the alterations in cellular factors responsible for a misshaped nucleus, which can then be mapped to a disease. This can be done using mechanical models that can decompose nuclear morphology into its contributing cellular factors. The theoretical models developed previously have simulated the changes in nuclear morphology from alterations in cellular factors such as osmotic pressure difference across the nuclear envelope (9), microtubules (9), and cell spreading (9, 11), and extracellular factors such as constrictions (12) and geometric constraints on cell shape (13). In contrast, here we present a mechanical model for quantitatively predicting the tension in actin cytoskeleton from the nucleus shape.

We modelled the nuclear envelope as an axisymmetric membrane compressed by tensed cortical actin (14) and inflated by a pressure arising from the osmotic pressure difference between the nucleoplasm and cytoplasm, microtubules and chromatin (10). Our model is akin to a tensegrity (15), which attains stability through tensions in cortical actin and the inflated nuclear envelope. Since cells are mostly flat, with much larger cell area in comparison to nuclear area, we simplify the force from cortical actin to that from a flat rigid plate (Fig. 1C) (9). By using the mechanical analysis developed in (16), we obtained two sets of nondimensional parameters that can describe nuclear morphology,

i. *η*_1_ = *PR*/2*E*_1_*H*, the ratio between the expanding pressure, P, to the elastic modulus, E_1_, of the nuclear envelope of radius, R, and thickness, H, in the unloaded state, and 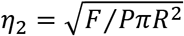, the ratio between the compressive force, F, to P
ii. *λ*_0_ (stretch indicator), the elastic stretch at the apex of the nucleus, and τ (flatness indicator), half of the angle subtended by cortical actin on the nuclear envelope in the undeformed state (Fig, 1B).

**Figure 1:**
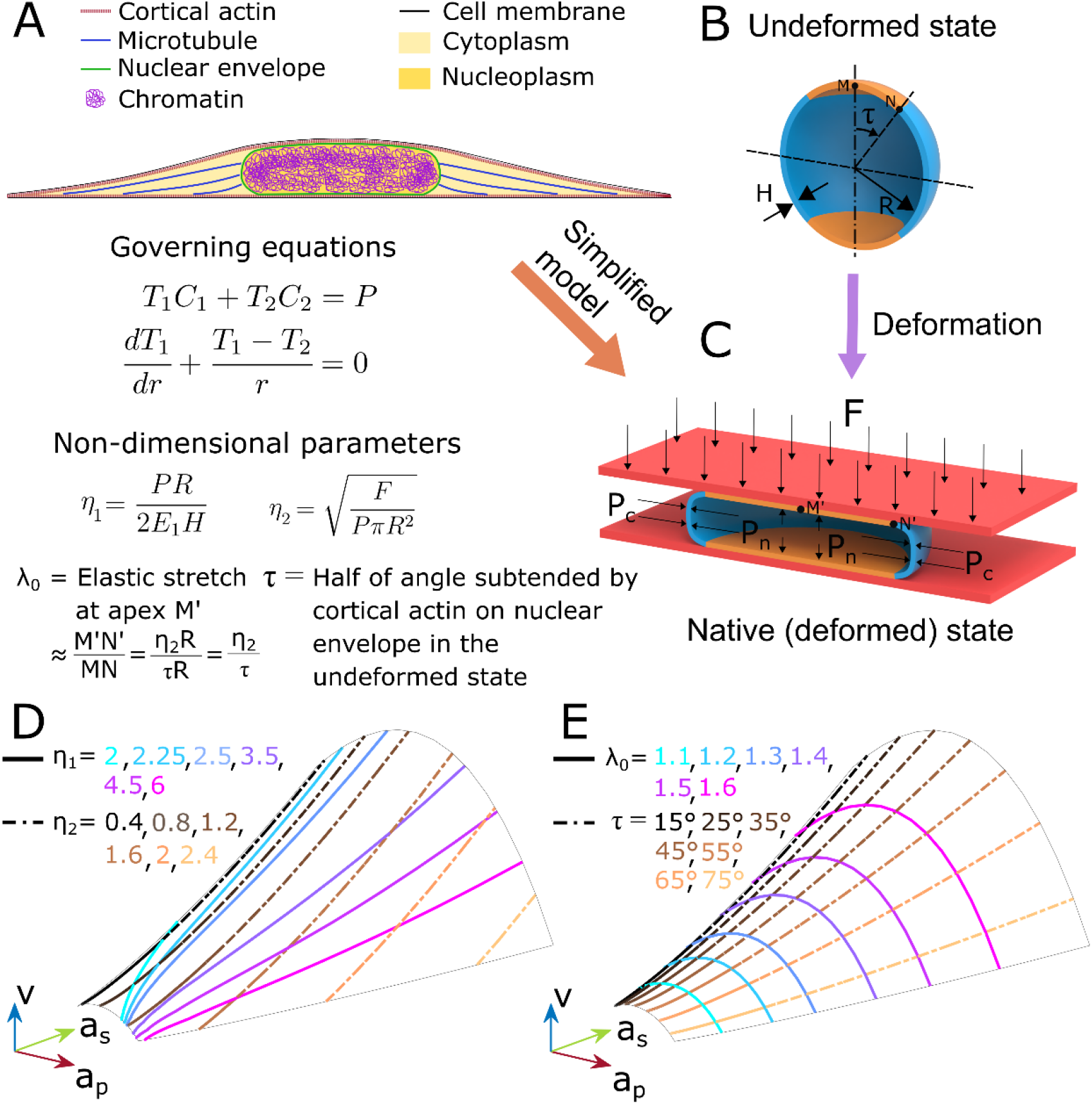
Two-parameter nondimensional model for nucleus morphology. (A) Nuclear envelope (blue) is shaped by forces from cortical actin (orange), microtubules (green), chromatin (purple), and an osmotic pressure difference between the nucleoplasm and cytoplasm. (B) The reference configuration, undeformed state, of the nuclear envelope is assumed to be a spherical membrane of radius *R* and thickness *H* (C) Forces from osmotic pressure, microtubules, and chromatin is lumped into an inflating pressure *P*=*P*_*n*_−*P*_*c*_. The force due to cortical actin, *F*,is assumed to be akin to a rigid flat plate pushing down on the nucleus. The equations of equilibrium of the nuclear envelope in terms of principal tensions, *T*_*1*_ and *T*_*2*_, and principal curvatures, *C*_*1*_ and *C*_*2*_, are shown. Solutions to these equations depend on two nondimensional parameters. Two choices for these nondimensional parameters are (i) *η*_*1*_ and *η*_*2*_, and (ii) *λ*_*0*_ and *τ*. By simulating the model for various values of these nondimensional parameters, we obtained the corresponding nuclear shapes. Normalized projected area (*a*_*p*_), surface area (*a*_*s*_) and volume (*v*), estimated from these nuclear shapes formed a surface in the *a*_*p*_-*a*_*s*_-*v* space since there are only two independent nondimensional parameters governing them. The contour lines of the nondimensional parameters, *η*_*1*_ and *η*_*2*_ (D), and *λ*_*0*_ and *τ* (E), on this surface are shown.

These parameters can be obtained by fitting our model to three nuclear shape parameters namely, projected area, surface area, and volume, of individual nuclei. *η*_*1*_ and *λ*_*0*_ represent the effect of pressure, *P*, on nucleus morphology, which is to expand or scale the nucleus. The elastic modulus of the nuclear envelope, *E*_*1*_, resists this expansion and therefore increase in *E*_*1*_ reduces *η*_*1*_ and *λ*_*0*_ and vice versa. *η*_*2*_ and *τ* represent the effect of force from cortical actin, i.e., the flatness of the nucleus. *η*_*2*_ is the ratio between the radius of the contact region between cortical actin and nuclear envelope (M′N′ in Fig. 1C) to the radius of the undeformed state, *R*. *τ* is the size of the contact region mapped to the undeformed state. It may be noted that our model has only two independent parameters and hence the values of both sets can be estimated from the other (Fig. 1D). For instance, *λ*_0_ ≈ *M*′*N*′/*MN* = (*η*_2_*R*)/(*τR*) ⇒ *η*_2_ ≈ *λ*_0_*τ* where *τ* is in radians. Since there are only two independent nondimensional parameters, the model predicts a relationship among projected area, surface area and volume of the nucleus. This relationship is represented by a surface in the three-parameter space of the aforementioned geometric quantities (Fig. 1D and E). Previously, we have used this model to predict molecular mechanisms responsible for changes in nuclear mechanics due to Hepatitis C Virus by merely analyzing the changes in nuclear morphology (17). In that work, we had focused on predicting modulus of nuclear envelope and thereby the expression of lamin-A,C from nuclear shape. Here, we focus on actin cytoskeletal tension and show that it can be predicted from changes in nuclear shape for multiple cell lines and under a variety of perturbations.

We first analyzed nucleus morphology of individual cells from multiple lines and observed that their variability is primarily unidirectional. This suggests that nucleus shape predominantly depends on a single variable. By using our model, we show that this principal variable is *λ*_*0*_. Next, we observed that the nondimensional parameters represented the mechanical state of the cell and hence, we hypothesized that the mechanical state of the cell could be estimated from them. To test this, we perturbed the tension in actin cytoskeleton using three independent processes – (i) direct depolymerisation of actin filaments using Cytochalasin D, (ii) upregulation of Rho/ROCK pathway by depolymerizing microtubules using Nocodazole, and (iii) by culturing cells on polyacrylamide gels of varying elastic moduli. We found that in each case *τ* predicted the modulation in actin cytoskeletal tension and *λ*_*0*_ corresponded to the expected variation in the modulus of the nuclear envelope. Finally, we show that *τ* is equivalent to ration between height and diameter, of the nucleus. We use this geometrical insight to (i) compare and validate our results with many previous studies and (ii) derive a convenient method for estimating *τ* from nuclear area and volume using an analytical expression and a graph.

## Results

### Variability in nuclear shape

To quantify the variability in nuclear morphology we acquired confocal images of nuclei of five cell lines namely, Huh7, HeLa, NIH3T3, MDAMB231 and MCF7. Nuclear surfaces were obtained (Fig. S1) from these confocal images using an 3D-active-contour-based image processing algorithm that we had previously developed (17). From the nuclear surfaces, we calculated three nuclear shape measures, namely, projected area, surface area, and volume. Next, we estimated the normalized projected area (*a*_*p*_), surface area (*a*_*s*_), volume (*v*) using the radius of the undeformed state, *R* (see Materials and Methods). These normalized shape parameters were fit to our model to obtain the nondimensional parameters, *η*_*1*_, *η*_*2*_, *λ*_*0*_, and *τ*, of each nuclei. Our two-parameter model could fit the three nuclear shape parameters at less than 5% error in each of these parameters, for more than 85% of the nuclei imaged from all cells lines.

The nuclear shape parameters corresponding to individual nuclei of all cell lines lie on the surface in the *a*_*p*_-*a*_*s*_-*v* space predicted by the model (Fig. 2A). Furthermore, the shape parameters corresponding to nuclei from each cell line nearly lie on straight lines (Fig. 2A). To obtain the direction of this straight line for each cell line, we calculated the eigenvectors of the covariance matrix of the normalized projected area, surface area and volume (Fig. 2B). The largest and second-largest eigenvectors were aligned along *λ*_*0*_ and *τ* respectively (Fig. 2B). We confirmed this by calculating the Pearson’s correlation coefficient of λ_0_ and τ with the components of nuclear shape parameters along the eigenvectors, v_1_ and v_2_. λ_0_ and τ showed high correlation with v_1_ and v_2_ respectively, whereas λ_0_ and v_2_, and τ and v_1_ were uncorrelated for all cell lines (Table S1). We obtained the standard deviation along the principal directions (*σ*_*1*_, *σ*_*2*_ and *σ*_*3*_) by calculating the square root of the eigenvalues (Table 1). *σ*_*1*_/*σ*_*2*_ was equal to 9.7, 3.8, 6.2, 6.8 and 5.1 for Huh7, HeLa, NIH3T3, MDAMB231 and MCF7 cells respectively. The large value of this ratio shows that the variability in nuclear shapes is mostly along one direction, i.e. along *λ*_*0*_. Variability along *τ* was lower than 25% of the variability along *λ*_*0*_. Hence, nuclear morphology is a single-variable function of *λ*_*0*_. Nuclear morphology obtained from an independent study (18) also exhibited this univariate behavior (last row of Table 1). This was further confirmed by the linear correlation between the other set of nondimensional parameters, *η*_*1*_ and *η*_*2*_ (Fig. S2A). Pearson’s correlation coefficient between *η*_*1*_ and *η*_*2*_ was 0.69, 0.9, 0.94, 0.9 and 0.89 for Huh7, HeLa, NIH3T3, MDAMB231 and MCF7 cells respectively. In contrast, λ_0_ and τ were uncorrelated (Fig. S2B). Pearson’s correlation coefficient between *λ*_*0*_, and *τ* was −0.04, −0.26, 0.30, 0.17, and −0.11 for Huh7, HeLa, NIH3T3, MDAMB231 and MCF7 cells respectively.

**Table 1:** Principal variability in nuclear morphology. Variability of HeLa cell nuclei from an independent study (18) is given in the last row.

| Cell line | Principal variability |  |  |
| --- | --- | --- | --- |
| | $\sigma_1$ | $\sigma_2$ | $\sigma_3$ |
| Huh7 | 5.6 | 0.6 | 0.2 |
| HeLa | 5.7 | 1.5 | 0.3 |
| NIH3T3 | 5.6 | 0.9 | 0.2 |
| MDAMB231 | 8.1 | 1.2 | 0.4 |
| MCF7 | 4.6 | 0.9 | 0.2 |
| HeLa (18) | 3.7 | 0.4 | 0.05 |

**Figure 2:**
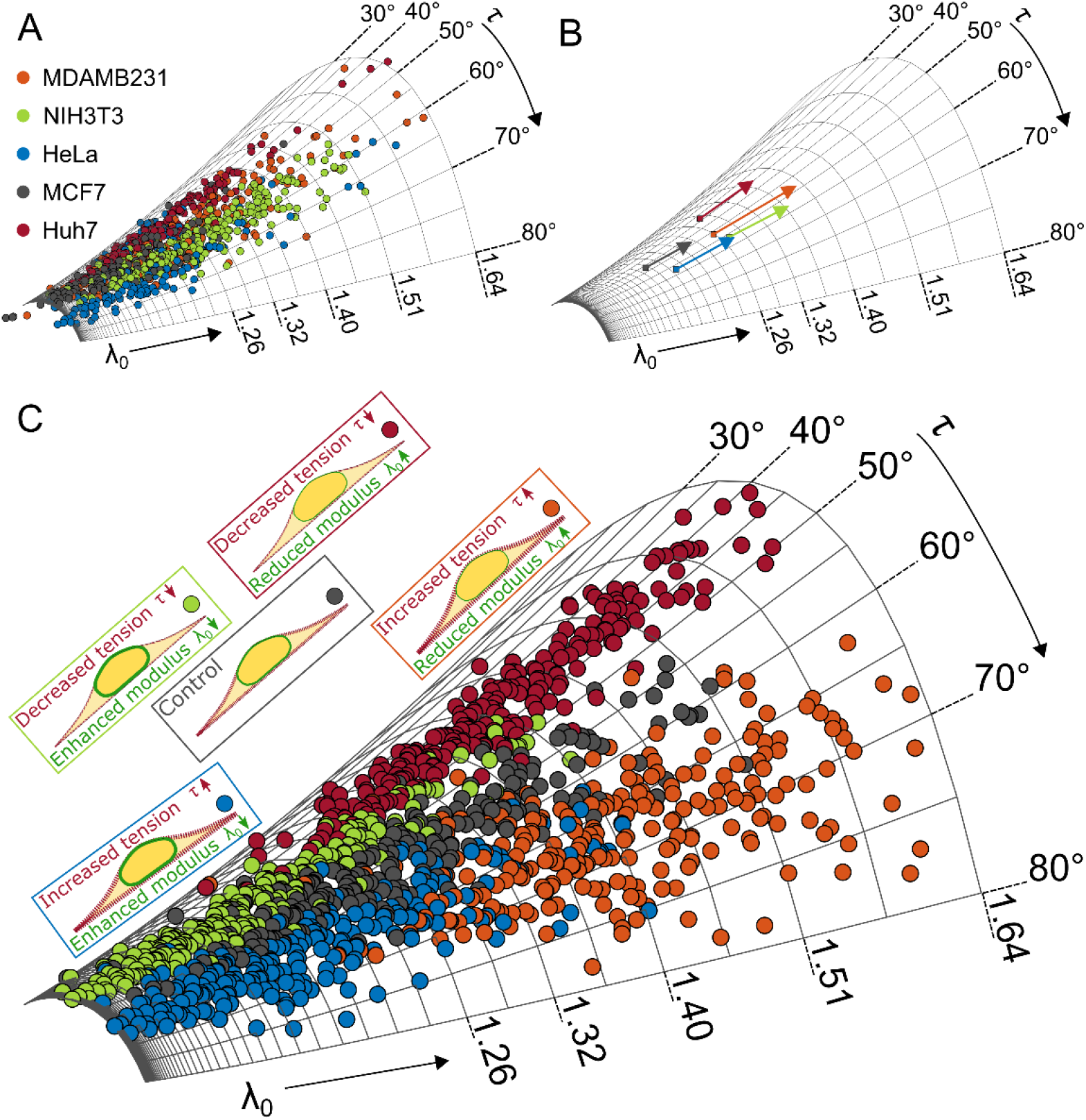
Variability in nucleus shape. (A) Nuclear shape parameters, normalized projected area, surface area and volume of individual nuclei of MDAMB231 (orange), HeLa (blue), MCF7 (grey), Huh7 (red) and NIH3T3 (green) cells plotted over the model surface (B) Direction of largest variability in nucleus shape was obtained by estimating the principal eigenvector of the covariance matrix of normalized projected area, surface area and volume of the nuclei. The length of the arrow is equal to the square root of the largest eigenvalue. We hypothesized that changes in actin cytoskeletal tension or elastic modulus of the nuclear envelope can be inferred from changes in *τ* and *λ*_*0*_ calculated from the nuclear morphology. It can also be qualitatively inferred visually from the location of nuclei on the model surface as shown in (C). The perturbations corresponding to different colors is shown beside the model surface. For example, decrease in tension and modulus will shift nuclei to lower *τ* and higher *λ*_*0*_ (red dots) in comparison to control (grey dots). It may be noted that the dots in (C) are for presenting the hypothesis and not actual nuclear shape data

Next, we checked if the range of values of *λ*_*0*_ and *τ* are a fundamental property of each cell line or whether they represent the mechanical state of the cells. For this we reduced the tension in actin cytoskeleton by culturing cells at a high cell density. *τ* was significantly lower in high-seeding-density cultures in comparison to those at low-seeding-density (Fig. S3) suggesting that these nondimensional physical parameters were not fundamental properties of the cells, but indicative of their mechanical state. Since *τ* represents the effect of force from cortical actin on nuclear envelope (Fig. 1) and *τ* decreased when actin cytoskeletal tension was reduced by increasing cell seeding density (Fig. S3), we hypothesized that *τ* could be used to predict actin cytoskeletal tension from nucleus shape. Furthermore, since stretch decreases with increase in elastic modulus of nuclear envelope, we hypothesized that *λ*_*0*_ could predict the changes in stiffness of the nuclear envelope. For testing these hypotheses, we systematically modulated the tension in actin cytoskeleton using three independent mechanisms – (i) downregulation by depolymerizing actin through Cytochalasin D treatment (ii) indirect upregulation by Nocodazole through activation of the Rho/ROCK pathway upon microtubule depolymerisation and (iii) mechanical modulation by growing on polyacrylamide gels of varying elastic modulus.

### Reducing the tension in actin cytoskeleton by treating with Cytochalasin D

We systematically reduced the tension in actin cytoskeleton by treating with increasing concentrations of Cytochalasin D, 0.46, and 0.92 μM, on four cell lines, HeLa, MCF7, MDAMB231, and Huh7. Nuclear morphology obtained from confocal images (Fig. S4) were fit to our model to estimate the nondimensional parameters of individual cells (Figs. 3 and S5). *τ* decreased and *λ*_*0*_ increased with increasing concentration of Cytochalasin D. Change in *τ* was analogous to the change in actin cytoskeletal tension. Increase in λ_0_ suggests either increase in the inflating pressure or decrease in the modulus of the nuclear envelope. Decrease in actin cytoskeletal tension has been shown to reduce lamin-A,C expression, and thereby reduce the modulus of the nuclear envelope (13, 19). Therefore, *λ*_*0*_ could be representing the modulus of nuclear envelope and lamin-A,C expression.

**Figure 3:**
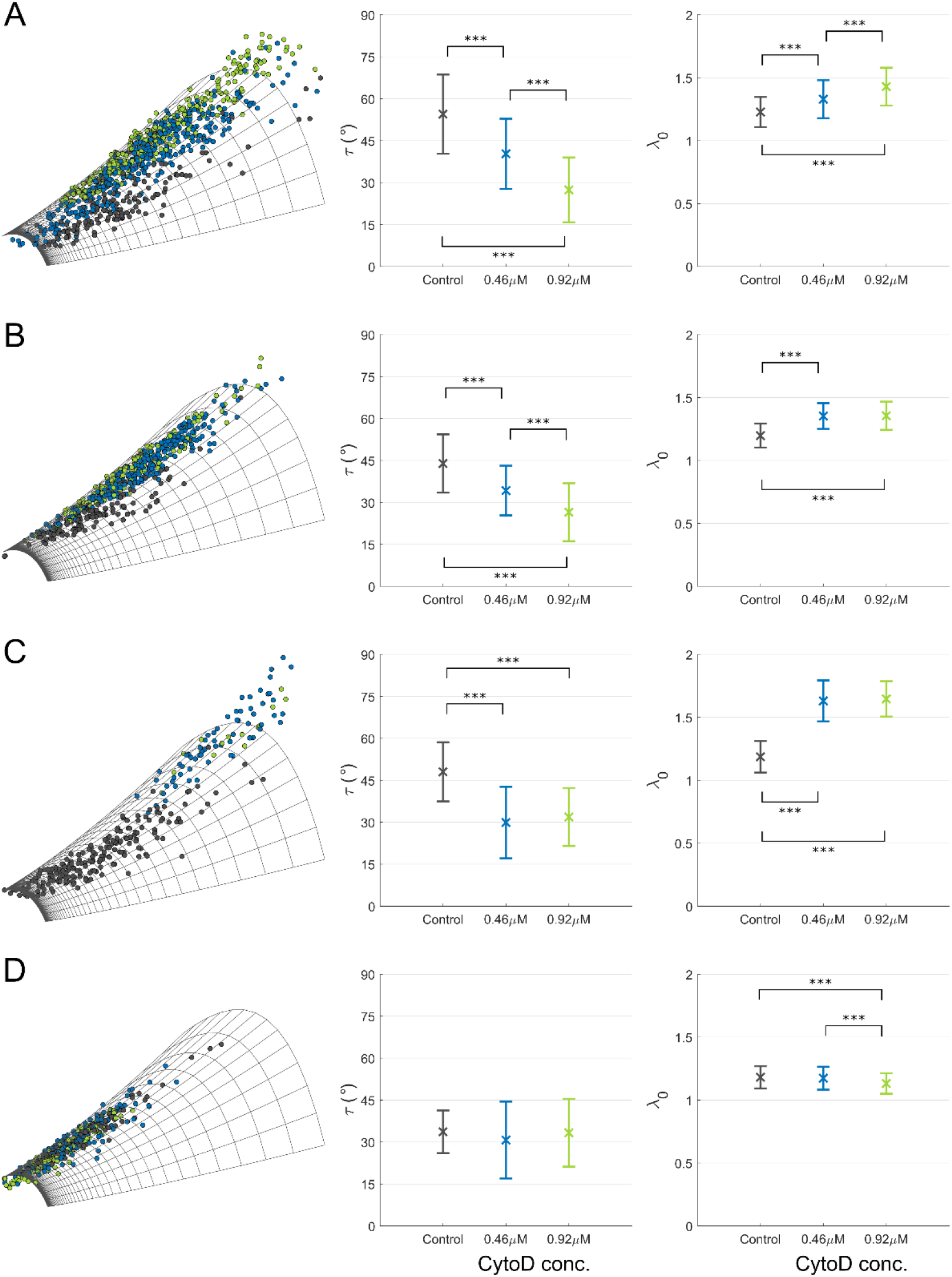
Changes in nondimensional parameters due to Cytochalasin D treatment. Four cell lines, HeLa (A), MCF7 (B), MDAMB231 (C), and Huh7 (D) were treated with two concentrations, 0.46 μM, and 0.92 μM of Cytochalasin D. By fitting our model to individual nuclei, nondimensional parameters *λ*_*0*_ and *τ* were estimated. Scatter plot of nuclear shape parameters on the model surface is shown in the left column. Each dot is an individual nucleus and the colors represent black – control, blue - 0.46 μM, and green - 0.92 μM of Cytochalasin D. Bar graphs with the mean and standard deviation of *τ* and *λ*_*0*_ are shown in the center and right columns respectively. Statistical analysis was performed using ANOVA with Bonferroni correction and significance levels are represented by * p<0.05, ** p<0.01, and *** p<0.001

In Huh7 cells, the decrease in τ was not significant (Fig. 3D). We hypothesized that this could be because of the low tension, *τ* ≈ 30°, in Huh7 control cells. Since the initial tension is low, further reduction in tension by Cytochalasin D treatment might not be detected. The nucleus is approximately spherical at low tension. Consequently, our assumption of treating the force from cortical actin like a rigid flat plate may not be valid. To test this, we treated cells with low actin cytoskeletal tension with Cytochalasin D. To obtain control cells with low actin cytoskeletal tension, we cultured them at high seeding density. We observed that at low tension, *τ* did not change consistently with Cytochalasin D treatment (Fig. S6). This suggests that there is a lower limit of detection for this technique of *τ* ≈ 30°. If the tension in control cells is lower than this limit, a further reduction in tension cannot be detected.

### Increasing the tension in actin cytoskeleton by treating with Nocodazole

Next, we increased the actin cytoskeletal tension by treating cells with Nocodazole. Depolymerizing microtubules by Nocodazole has been shown to increase the tension in actin cytoskeleton (20, 21) via the Rho/ROCK pathway (22–25). Huh7, MCF7, MDAMB231 and NIH3T3 cells were incubated with 6 μM Nocodazole for 2 h. From confocal images of the nuclei (Fig. S7), we obtained the nondimensional parameters by fitting our model. We observed that *τ* increased significantly in all the cell lines whereas the variation in *λ*_*0*_ was lower and inconsistent across cell lines (Figs. 4 and S8). The consistent increase in *τ* across cell lines shows that it can predict changes in actin cytoskeletal tension even when modulated using an indirect signaling mechanism. In case of *λ*_*0*_, the disruption of microtubules abrogates the compressive pressure exerted by them, thereby increasing *P* and *λ*_*0*_. On the other hand, enhanced tension in actin cytoskeleton is known to increase elastic modulus of the nuclear envelope, *E*_*1*_, by upregulating lamin-A,C (13, 19), thereby decreasing *λ*_*0*_. These competing mechanisms could have balanced each other and stabilized *λ*_*0*_.

**Figure 4:**
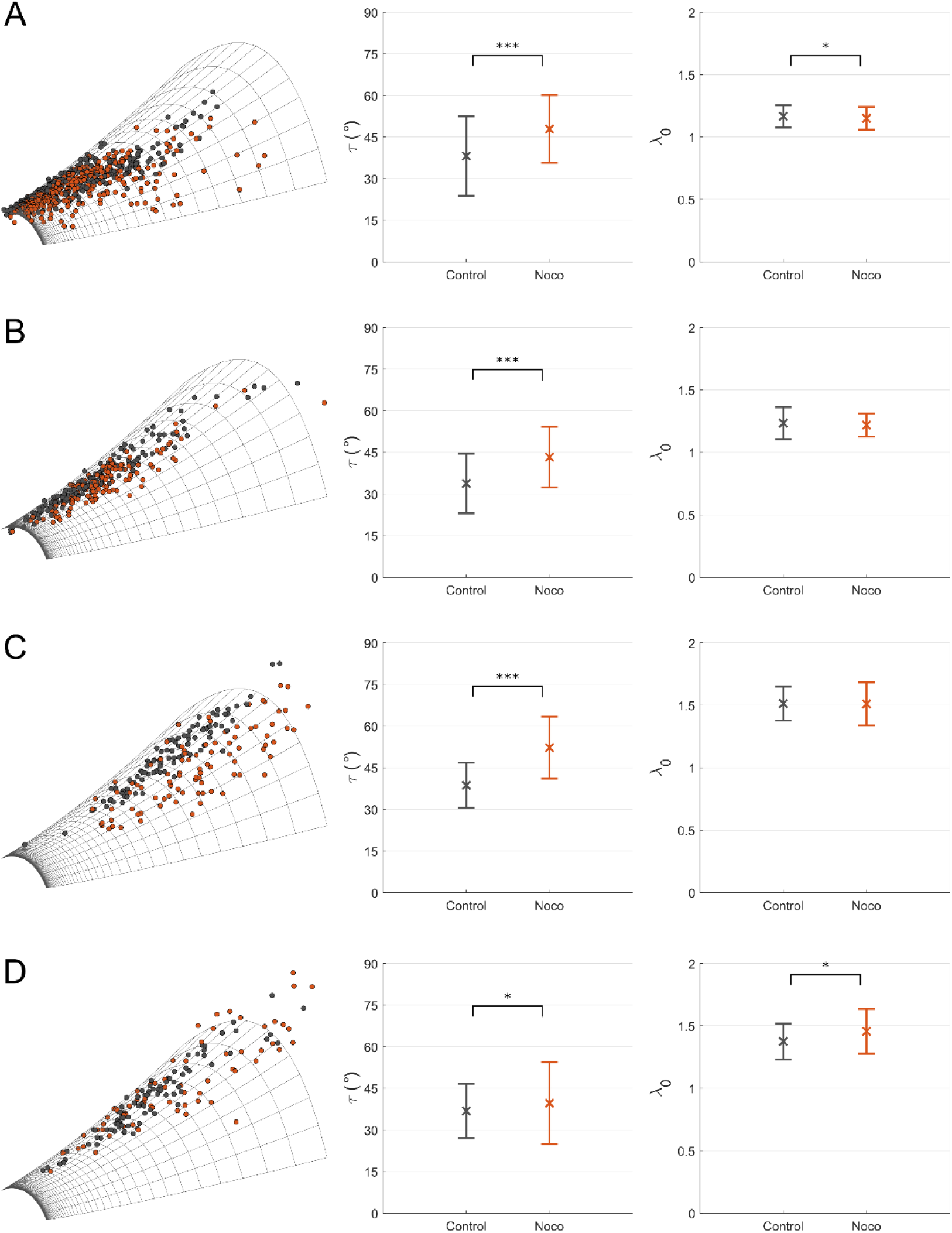
Changes in nondimensional parameters due to Nocodazole treatment. Four cell lines, (A) Huh7, (B) MCF7, (C) MDAMB231, and (D) NIH3T3, were treated with 6 μM Nocodazole. By fitting our model to individual nuclei, nondimensional parameters *λ*_*0*_ and *τ* were estimated. Scatter plot of nuclear shape parameters on the model surface is shown in the left column. Each dot is an individual nucleus and the colors represent black – control, and red – Nocodazole-treated cells. Mean and standard deviation of *τ* and *λ*_*0*_ are shown in the center and right columns respectively. Statistical analysis was performed using Kolmogrov-Smirnov test and significance levels are represented by * p<0.05, ** p<0.01, and *** p<0.001

### Modulating the tension in actin cytoskeleton by varying substrate stiffness

We further tested if our method can predict the modulation in actin cytoskeletal tension due to passive mechanical signals. For this, we cultured Huh7 cells on polyacrylamide gels of varying stiffness. Previously, hMSCs cultured on polyacrylamide gels of increasing elastic modulus were shown to have enhanced actin cytoskeletal tension and lamin-A,C expression (19). We fabricated polyacrylamide gels of 2.5 kPa, 11 kPa, and 36 kPa elastic modulus. Glass coverslip of elastic modulus ≈ 1 GPa was used as control. The nondimensional parameters were obtained by fitting our model to the morphology of individual nuclei obtained from confocal images (Fig. S9). We observed that *τ* increased and *λ*_*0*_ decreased with increasing substrate elastic modulus (Figs. 5 and S9). This corroborates with previous experimental results on enhanced cytoskeletal tension and lamin-A,C expression with increasing substrate stiffness (19).

**Figure 5:**
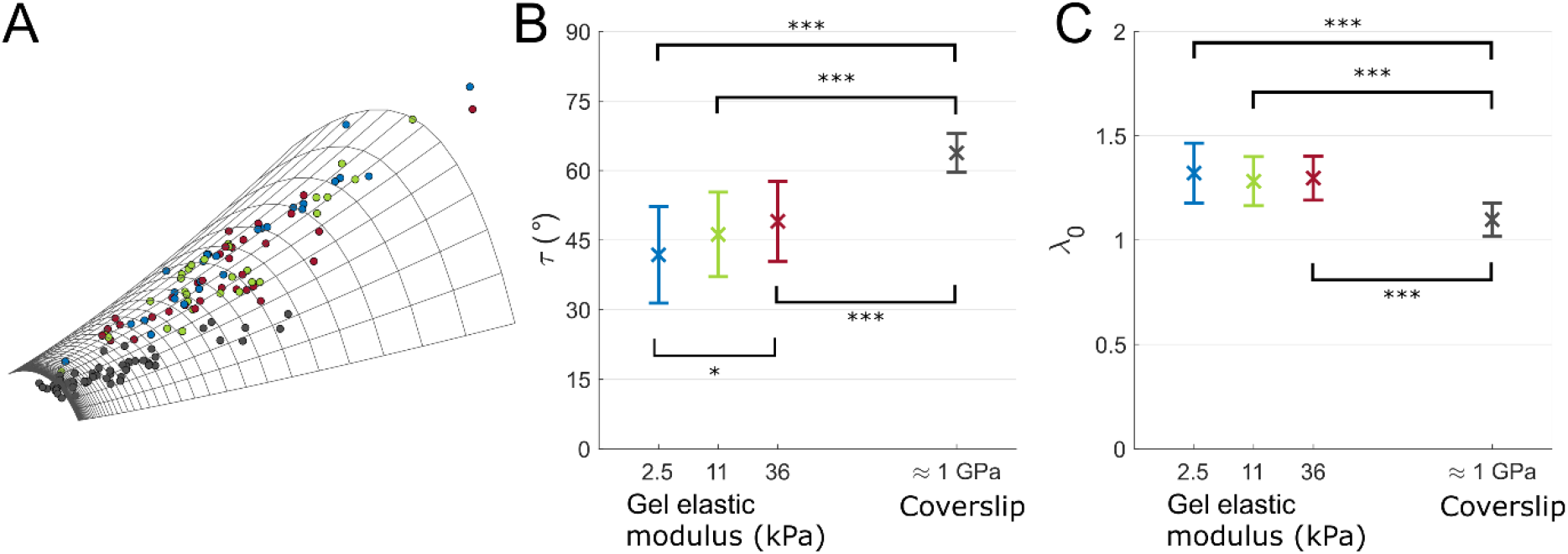
Changes in nondimensional parameters on varying elastic modulus of substrate. Huh7 cells were cultured on polyacrylamide gels with elastic modulus of 2.5, 11, and 36 kPa. By fitting our model to individual nuclei, nondimensional parameters *λ*_*0*_ and *τ* were estimated. (A) Scatter plot of nuclear shape parameters on the model surface. Each dot is an individual nucleus and the colors represent black – control (coverslip ≈ 1 GPa), blue – 2.5 kPa, green – 11 kPa and red – 36 kPa. Mean and standard deviation of *τ* and *λ*_*0*_ are shown in (B) and (C) respectively. Statistical analysis was performed using ANOVA with Bonferroni correction and significance levels are represented by * p<0.05, ** p<0.01, and *** p<0.001

### Geometric interpretation of nondimensional parameter

We observed that the aspect ratio, height to diameter, of the nuclear geometry simulated by our model is independent of *λ*_*0*_ (Fig.6A). Aspect ratio decreased slightly for *λ*_*0*_ < 1.2 and remained constant for *λ*_*0*_ > 1.2 for all values of *τ* (Fig. 6A). Aspect ratio decreased with *τ* for all values of *λ*_*0*_ (Fig. 6A). Therefore, aspect ratio is an equivalent measure to *τ*. To test this, we calculated the correlation between aspect ratio and the component of the nuclear shape parameters along the principal directions of variability for the cells shown in Fig. 1 and Table S1. Aspect ratio was negatively correlated with the second principal direction, Pearson’s correlation coefficient for aspect ratio was −0.88, −0.94, −0.83, −0.8, −0.94 for Huh7, HeLa, NIH3T3, MDAMB231, and MCF7 cells respectively. Furthermore, aspect ratio was uncorrelated with the first principal direction. Pearson’s correlation coefficient was −0.3, −0.11, −0.43, −0.42, −0.1 for Huh7, HeLa, NIH3T3, MDAMB231, and MCF7 cells respectively. Hence, aspect ratio could be used as a predictor for actin cytoskeletal tension. This makes sense intuitively because higher tension flattens the cells thereby decreasing the aspect ratio of the nucleus. We used this geometrical insight into *τ* for comparing our results with previous studies and for deriving a convenient relation to estimate the aspect ratio, and thereby *τ*, from nuclear volume and projected area.

**Figure 6:**
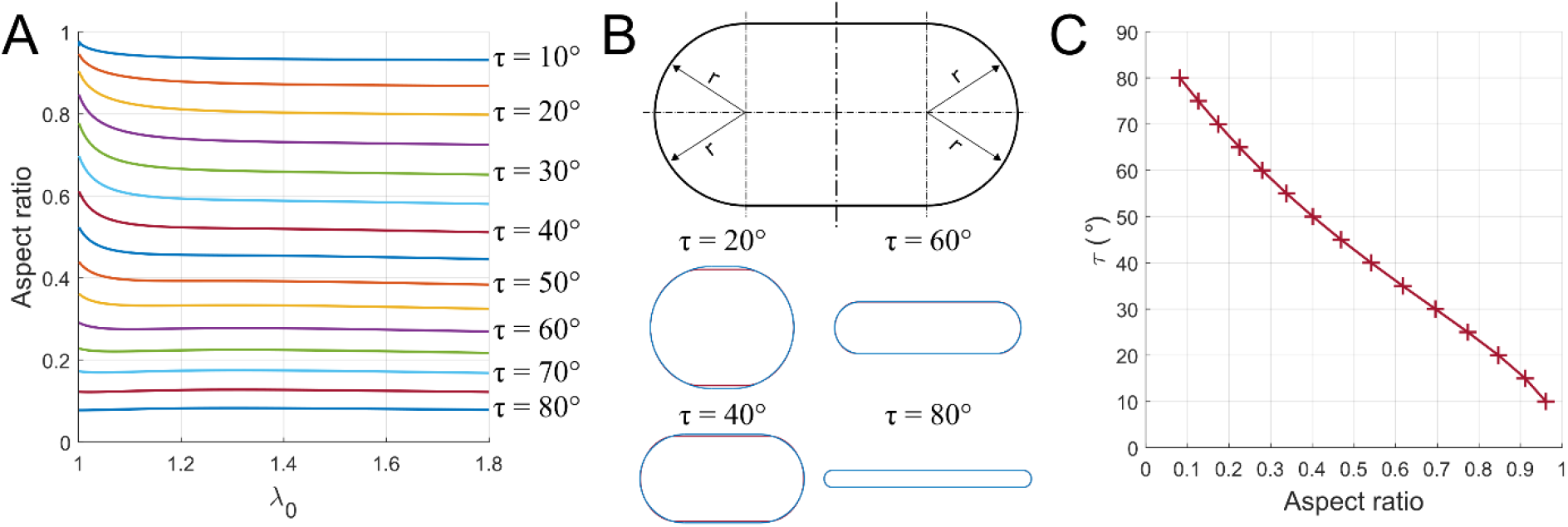
Geometric approximation of the model. (A) Variation of aspect ratio, height to diameter, of simulated nucleus shape with *λ*_*0*_ for different values of *τ*. (B) Approximate geometry of the nucleus, axisymmetric surface with cross-section of a rectangle with semi-circular ends. Comparison of approximation (blue) with simulated nuclei (red) for *λ*_*0*_ = 1.2 and τ = 20°, 40°, 60°, and 80°. (C) *τ* for different values of aspect ratio obtained by fitting the approximate geometry to nuclear volume and projected area

Many studies have reported changes in nuclear height and projected area as a consequence of perturbations to cellular and extracellular factors governing nuclear shape. From these measurements, alternation in aspect ratio can be inferred, which can be used to compare the predictions of our model with their experimental observations.

i. Aspect ratio of nuclei increased when cells were treated with tension-reducing agents such as Latrunculin A (Fig. 1e in (6)), blebbistatin (Fig. 3d in (26)), and Y-27632 (Fig. SE10 in (27)).
ii. Aspect ratio of nuclei decreased when *Drosophilla* S2R+ cells were treated with Nocodazole (Fig. 1e in (6)).
iii. Aspect ratio of nuclei of hMSCs (Fig. 1 in (19)) and NIH3T3 (Fig. 1 in (28)) cells decreased when grown on substrates of increasing elastic moduli.
iv. Aspect ratio of nuclei increased when microtubules were stabilized by treating with Paclitaxel (Fig. 1e in (6)). Our model predicts that this could be due to decrease in actin cytoskeletal tension. This prediction has been confirmed by multiple independent studies, which have shown that Paclitaxel treatment decreases cellular traction forces (29, 30).

All these studies independently validate the predictions from our model.

Our results show that irrespective of the biochemical mechanism used to modulate the tension in actin cytoskeleton or cell line, the modulation can be inferred from the nucleus shape using our model. This suggests that the connection between the actin cytoskeletal tension and nucleus shape is essentially mechanical.

The emerging mechanical picture for a cell is akin to a tent; stabilized by the tension in cortical actin (similar to the canopy), which is supported by the nucleus (akin to the pole in the middle that carries compression). The nuclear envelope acquires compression-carrying capacity through a net inflating pressure that tenses it, akin to a balloon. Since the structure is stabilized by the tensions in cortical actin and nuclear envelope, this mechanical viewpoint is akin to a tensegrity with the following distinctions from the original tensegrity theory for cells (31) - (i) the tensile element is a two-dimensional membrane (ii) primary compression-carrying member is the nucleus, which is an inflated membrane, and therefore a tensegrity by itself. Our study shows that this simple mechanical picture not only satisfies multiple experimental observations, but also enables prediction of the changes in one member by measuring the changes in the other.

We further derived a convenient technique to apply our model for predicting actin cytoskeletal tension from nucleus shape. We observed that the simulated nuclear shapes can be approximated by a flat ‘pancake’ geometry with circular ends (Fig. 6B). For such geometries, the aspect ratio, *γ*, can be shown to be the solution of the cubic equation,

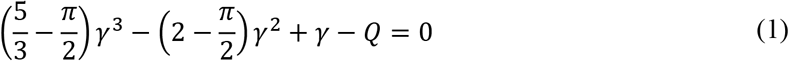

where *Q* is a nondimensional function of nuclear volume *V* and projected area *A* given by 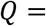 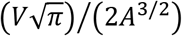 (see Supplementary Information for derivation). Analytical expression for the solution of this cubic equation is

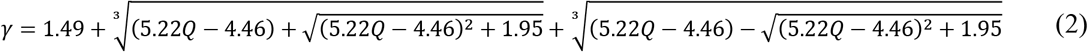

Since *γ* is independent of *λ*_*0*_ and a function of only *τ* (Fig. 6A), *τ* can be obtained from *γ* (Fig. 6C). It may be noted that *Q* is similar to Vogel number, which is a measure of the flatness of an object. Vogel number is defined as the ratio of the square root of the surface area to the cube root of the volume (32). *Q* is therefore proportional to the inverse of the cube of Vogel number.

We summarize the following techniques of increasing accuracy and complexity for estimating τ from nucleus shape

i. From nuclear volume, *V*, and projected area, *A*, calculate 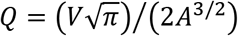. Obtain the aspect ratio, *γ*, from *Q* by solving Eq. (1) or by using the expression in Eq. (2). Now, estimate *τ* from *γ* using the graph in Fig. 6C. The points used to plot the graph in Fig. 6C are given in the excel file ‘aspectratio_tau.xls’ in Supplementary Data.
ii. Assume a suitable value for *R* and estimate the normalized projected area, surface area and volume of nuclei (See materials and methods). Plot the normalized nuclear shape parameters on the model surface and obtain the nondimensional parameters corresponding to the nearest grid point. MATLAB scripts and data files for performing this fit is given in the Supplementary Data.
iii. Numerically fit the model by minimizing the error in normalized projected area, surface area and volume between the simulated and experimentally measured nuclei (See materials and methods).

Since our model has only two independent parameters, two nuclear shape measures are sufficient to fit the model. Additional shape measures, such as surface area, ensure a better fit by correcting for any errors in estimation of nuclear shape measures. Hence, the method using only two nuclear shape measures, volume and projected area, should be employed only when the images are of high quality. We recommend signal to noise ratio > 2 for this simplified method. We have derived this method using nuclear volume and projected area because these shape measures can be easily obtained from confocal images by thresholding.

## Discussion

We have shown that by using a nondimensional mechanical model, the different contributions to the nucleus shape can be separated into their individual contributions. We modulated the tension in actin cytoskeleton using three independent mechanisms and showed that the modulation in each case could be predicted by *τ*, half of the angle subtended by the actin cap on the nuclear envelope in the undeformed state (Figs. 3-5). Changes in the elastic modulus of nuclear envelope could be predicted from *λ*_*0*_, stretch in the nuclear envelope at its apex (Figs. 3-5). We further showed that *τ* is equivalent to the aspect ratio of the nucleus, which enabled us to compare and validate our method with many previous experimental observations (6, 26–28).

Our method can be used to check the effect of various perturbations such as chemical, biological and mechanical on the actin cytoskeletal tension by merely analyzing nucleus shape. In this work we have shown this for two chemical perturbations, Cytochalasin D (Fig. 3) and Nocodazole (Fig, 4), and a mechanical perturbation, varying substrate stiffness (Fig. 5). Previously, we have shown this for a biological perturbation, Hepatitis C Virus in liver cells (17). Hence, our method can be used as convenient alternative to other methods for estimating actin cytoskeletal tension (33) through cell contractility such as traction force microscopy using microbeads (34) and micropillars (35), and gel contraction assays (36). Many cellular processes are regulated by the tension in actin cytoskeleton and hence alterations in it are associated with many diseases such as asthma (37), hypertension (38) and cancer (39). For example, different cancer states such as epithelial, hybrid and mesenchymal have varying actin cytoskeletal tension (40). Since our method calculates *τ* for individual cells from the shape of its nucleus, we could estimate the actin cytoskeletal tension for individual cells in a tumor. This can potentially identify the cancer state of each cell in a tumor and hence could be a technique to quantify cancer heterogeneity. Therefore, our technique could be an experimental and diagnostic tool.

Previously we have used nondimensional parameters *η*_*1*_ and *η*_*2*_ to predict the changes in the expression of cytoskeletal and nuclear envelope proteins by Hepatitis C Virus by merely analyzing the changes in nuclear shape (17). Here we found that these two nondimensional parameters are positively correlated under all perturbations to actin cytoskeletal tension done in this study (Figs. S5, S8 and S10). This could be because of molecular mechanisms interrelating osmotic pressure and actin cytoskeletal tension (41). In contrast, the other set of nondimensional parameters, *λ*_*0*_ and *τ* correspond to the principal directions of variability and are independent of each other. This independence can be inferred from the orthogonality of the contour lines for *λ*_*0*_ and *τ* in Fig. 1D. *λ*_*0*_ and *τ* estimated from the nuclei of a cell population are uncorrelated (Fig. S2B), further confirming that these parameters are independent. Another important feature of *τ* is that it is independent of *R*, the radius of the nucleus in the reference configuration (Fig. 1B). This is a very important property because the undeformed configuration is unobservable. Having to assume this unobservable undeformed state is a fundamental drawback of biomechanics studies that employ a solid mechanics approach. The independence with the initial configuration allows us to define a lower limit, *τ* > 30°, for detecting a reduction in actin cytoskeletal tension. Even though *η*_*2*_ varies analogous to *τ* (compare Figs. 3-5 with Figs. S5, S8 and S10), we cannot define a lower limit of detection for *η*_*2*_ because it depends on the value chosen for *R*. Therefore, we recommend using τ for predicting actin cytoskeletal tension from nuclear shape.

A limitation of our method is that decrease in actin cytoskeletal tension cannot be detected if the control cells have low tension. We have estimated this lower detection limit as, *τ* ≤ 30°. This restriction is because we have simplified the force from cortical actin to compression from a rigid flat plate. At low tension, this assumption may not be valid. Since we are observing only nuclear shape (dimension = length), our nondimensional parameters derived from nuclear geometry reflect the ratio of force (dimension = Newtons) to stiffness (dimension = Newtons per meter). For instance, *λ*_*0*_ and *η*_*1*_ correspond to ratio between inflating pressure and elastic modulus. Therefore these parameters may not be able to distinguish between an increase in inflating pressure and a decrease in elastic modulus. It is also pertinent to note another interpretation for nuclear mechanics, which assumes that changes in nuclear size are due to unfolding of wrinkles in the nuclear envelope and not due to its stretching (42, 43). These authors have also reported that aspect ratio of the nucleus decreases and do not change with myosin inhibitors ML7 and Y-27632 respectively (11).

We showed that nucleus morphology in a cell population is primarily a single-variable function of *λ*_*0*_. The variability in *τ* accounts for less than 25% of the total variability in nuclear shape. This is interesting because there are multiple biomechanical and biochemical parameters that govern nuclear shape. In our model we have considered the following biomechanical parameters - tension in cortical actin, elastic modulus of the nuclear envelope and osmotic pressure difference between the nucleoplasm and cytoplasm, which are dependent on the following biochemical parameters - expression of actin, myosin, and lamin. The univariate behavior of nucleus shape suggests that these biomechanical and biochemical parameters collapse into a single parameter through interdependent signaling mechanisms. For example, lamin-A,C is known to regulate myosin through the SRF pathway (19). Furthermore, since the principal variability is along *λ*_*0*_, which is equivalent to scaling, this variability could be cell cycle dependent. This is because cell cycle progression is known to increase nuclear size to accommodate the increasing DNA content (44). A cell-cycle-dependent mechanism can also explain the variability in *τ* because traction forces, and therefore the actin cytoskeletal tension, is known to vary with cell cycle (45). Therefore, the variability in nuclear shapes could be arising out of a cell-cycle-dependent mechanism with biochemical feedback to integrate all the independent variables to a single variable.

In summary, we have used a mechanical model to decompose the contributions from actin cytoskeletal tension and elastic modulus of the nuclear envelope to nuclear shape. The nondimensional parameters *τ* and *λ*_*0*_ that correspond to these physical properties of the cell, were shown to be the principal variables along which nuclear shape varies in a cell population. By validating our model with multiple perturbations across many cell lines and previous studies, we propose a mechanical picture of the cell akin to a tent, which is stabilized by the tensions in actin cytoskeleton and nuclear envelope. We further derive a convenient method to estimate *τ* from nuclear volume and projected area and thereby predict the actin cytoskeletal tension. From a larger perspective, our results show that by combining physical principles with experimental observations deeper physical insights can be derived.

## Materials and methods

### Cell culture, chemical treatments and immunofluorescence

All cell lines were cultured at 37 °C in DMEM with 10% FBS. Cells were regularly passaged at around 80% confluence. For the Cytochalasin D and Nocodazole treatments, cells were seeded at low concentrations (around 100 k cells on a 22 mm circular coverslip) and allowed to attach overnight. 16 h after seeding, the cell medium was replaced with another containing the chemical at the required concentration for 2 h. For our studies we have used 0.46 μM, 0.92 μM and 1.8 μM solutions of Cytochalasin D and 6 μM solution of Nocodazole in DMEM. After incubation for 2 h, the cells were fixed using 4% paraformaldehyde and stained for nucleus and actin using Hoechst 33342 and Rhodamine Phalloidin respectively. Confocal z stacks of the stained cells were taken on a Leica Microsystems, TCS SP5 II confocal microscope. An oil-immersion objective lens with a magnification of 63X and a numerical aperture of 1.4 was used. Z stack images were taken at a pixel size of 240 nm in the lateral directions and z-step size of 500 nm. The morphology of the nucleus was obtained from these confocal stacks using a 3D-active-contour-based image processing technique developed previously (17).

### Polyacrylamide gel fabrication and characterization

We have followed the protocol for polyacrylamide gel fabrication reported in (46). Coverslips were ultrasonicated in 10% Extran solution for 15 mins and kept in hot-air oven at 80°C for 30 mins. The coverslips were then washed with DI water and the traces of water were removed by rinsing with 100% ethanol. The cleaned coverslips were then kept overnight for drying at 80°C in the hot-air oven. For better adhesion to gels, these coverslips were activated by treating with 10% 3-(aminopropyl)-triethoxysilane for 15 mins and subsequently washed with DI water and treated with 0.5% glutaraldehyde for 30 mins. The coverslips were washed again and dried in the laminar air-flow hood and sterilized using UV.

We fabricated three bioinert polyacrylamide gels of different elastic moduli, low, intermediate and high, by altering the relative concentrations of acrylamide monomer and bis-acrylamide cross-linking monomer. Polyacrylamide gel precursors were prepared by mixing 10% v/v acrylamide (AA, 40%, Sigma) with 0.03%, 0.1%, and 0.3% v/v of N,N’-methylenebisacrylamide (bis-AA 2% w/v in DW, Sigma) in DI water for low, intermediate and high elastic modulus gels respectively. Gelation was initiated by adding 0.1% v/v tetramethylethylenediamine (TEMED, Sigma) and 0.1% w/v ammonium persulfate (Sigma) to the gel precursor. 30 μL of this solution was pipetted to silanized coverslips, covered with extran-treated glass slides and cured for 30 mins. The coverslip with the intact gel was carefully peeled off from the glass slide. Next, we treated these gels with Sulpho-SANPAH to enable cell adhesion. Gels were immersed in 1 mg/mL sulfo-SANPAH (Fisher Scientific) in 50 mM, pH 8.5 HEPES and reacted under UV for 10 min.

The treated gels were washed thrice with HEPES buffer. To improve cell adhesion, we coated the gels with collagen. Rat tail collagen - I was mixed in 0.1 M acetic acid (Fisher Scientific) at equal volume and in 50mM, pH 8.5 HEPES to reach 0.1 mg/ml final concentration. Gels were immersed in this collagen solution and incubated overnight at 4 °C. Prior to cell seeding, the gels were sterilized by UV inside the laminar air-flow hood for 20 mins. The gels were maintained in a hydrated state all throughout these steps.

The elastic modulus of the gels were determined using an Atomic Force Microscope (XE Bio from Park Systems, Suwon, South Korea). We have used a V-shaped cantilever with a spherical bead of diameter 5.2 μm made of silicon dioxide attached to its bottom (AppNano HYDRA6V-200NG-TL; AppNano, Mountain View, CA). The force-displacement curves were fit to a Hertzian contact model to determine the elastic modulus (17). We obtained the following elastic moduli for the gels: low – 2.5 ± 0.3 kPa, intermediate - 10.7 ± 0.2 kPa, and high - 36 ± 2.2 kPa.

### Nondimensional mechanical model for the nucleus

We assumed that the nucleus is shaped by two forces, an expanding pressure, *P* and a downward compressive force *F* (9). The inflating pressure is the net result of the osmotic pressure difference between the nucleoplasm (*P*_*n*_) and cytoplasm (*P*_*c*_), compressive pressure from the microtubules, and chromatin (12). The compressive force is the downward force due to cortical actin and was assumed to be akin to a flat rigid plate pushing down on the nuclear envelope. The nuclear envelope was assumed to be a hypereslastic membrane (incompressible Mooney-Rivlin material) that is spherical in the unloaded state (9, 12, 47).

Since the initial geometry, forces and boundary conditions are axisymmetric, we used an analytical formulation developed for mechanical equilibrium of axisymmetric membranes (16). The solution to the governing equations for these boundary conditions depend only on two nondimensional parameters, (i) *η*_1_ = *PR*/2*E*_1_*H*) and (ii) 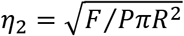 where *P* is the expanding pressure, *E_1_* is the elastic modulus of the nuclear envelope, *F* is the compressive force from cortical actin, *R* is the radius and *H* is the thickness of the nuclear envelope in the undeformed state. *η_1_* appears in the governing equation, and *η_2_* comes from the boundary condition. For solving these differential equations, we require two other parameters: (i) *λ_0_*, the stretch at the apex point of the nuclear envelope (point M′ in Fig. 1C) and (ii) *τ*, half of the angle subtended by the contact region between cortical actin and nuclear envelope in the undeformed state (Fig. 1B). However, as there are only two independent parameters, by specifying either of them, the other two can be determined (17). In our simulations we have specified λ_0_ and τ, and estimated η_1_ and η_2_.

We first solved the forward problem, ie, to estimate the nucleus morphology for a given set of nondimensional parameters. For given *λ*_*0*_ and *τ*, we first calculated *η*_*1*_ and *η*_*2*_, and then numerically integrated the governing equations to obtain the normalized nuclear morphology, which is the deformed shape when a spherical membrane of unit radius, thickness and modulus of elasticity is deformed by an inflating pressure = *η*_*1*_ and compressive force = *η*_*2*_. The actual nuclear morphology can be obtained by scaling the normalized nuclear morphology by *R*. The actual and normalized nuclear shape parameters are related through the following scaling relations - *A*_*p*_ = *R*^2^*a*_*p*_, *A*_*s*_ = *R*^2^*a*_*s*_, and *V* = *R*^3^*v*, where *A*_*p*_, *A*_*s*_ and *V* are the actual projected area, surface area and volume of the nucleus and *a*_*p*_, *a*_*s*_ and *v* are the corresponding normalized quantities.

Next, we used this forward problem to fit our model and obtain nondimensional parameters corresponding to experimentally measured nuclei. We calculated the projected area, surface area and volume of individual nuclei from the nuclear surfaces obtained by applying our image processing method on the confocal images. These nuclear shape parameters were normalized using the following relations - 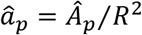, 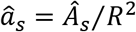, and 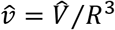 where ‘^’ indicates the experimentally measured quantities. The model was fitted to individual nuclei by minimizing the error between the simulated and experimentally measured normalized nuclear morphologies.

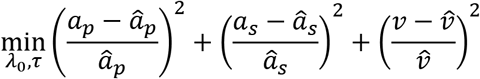

We have considered only those fits in which the error between the model and experimentally measured nuclei was less than 10% for each of the nuclear shape parameters. More than 90% of nuclei from each experiment were found to fit within this error threshold. Since we have neglected bending modulus in the nuclear envelope, it will buckle under compression and therefore *λ*_*0*_ > 1. Hence, *R* was chosen such that more than 90% of the nuclei satisfy this criteria on *λ*_*0*_. Nuclei with *λ*_*0*_<1 were not considered in our analysis. Furthermore for each experiment, we have used the same *R* for all the nuclei of a cell line since individual cells of each cell line were assumed to be descendant from a single clone. More details of the model is available in our previous publication (17).

## Acknowledgments

Authors would like to thank Reshma Pradeep and Suma M.S. for the preliminary experiments, and Sachin Arya for some of the illustrations

## References

1. J. Irianto, C. R. Pfeifer, I. L. Ivanovska, J. Swift, D. E. Discher, Nuclear Lamins in Cancer. Cell. Mol. Bioeng. 9, 258–267 (2016).

2. J. I. De las Heras, D. G. Batrakou, E. C. Schirmer, Cancer biology and the nuclear envelope: A convoluted relationship. Semin. Cancer Biol. 23, 125–137 (2013).

3. J. Lammerding, et al., Lamin A / C deficiency causes defective nuclear mechanics and mechanotransduction. J. Clin. Invest. 113, 370–378 (2004).

4. H. J. Worman, C. Ostlund, Y. Wang, Diseases of the nuclear envelope. Cold Spring Harb. Perspect. Biol. 2, 1–17 (2010).

5. E. S. Folker, C. Ostlund, G. W. G. Luxton, H. J. Worman, G. G. Gundersen, Lamin A variants that cause striated muscle disease are defective in anchoring transmembrane actin-associated nuclear lines for nuclear movement. Proc. Natl. Acad. Sci. 108, 131–136 (2011).

6. N. M. Ramdas, G. V. Shivashankar, Cytoskeletal Control of Nuclear Morphology and Chromatin Organization. J. Mol. Biol. 427, 695–706 (2015).

7. P. Scaffidi, T. Misteli, Lamin A-Dependent Nuclear Defects in Human Aging. Science (80-.). 312, 1059–1063 (2006).

8. S. M. Schreiner, P. K. Koo, Y. Zhao, S. G. J. Mochrie, M. C. King, The tethering of chromatin to the nuclear envelope supports nuclear mechanics. Nat. Commun. 6, 7159 (2015).

9. D.-H. Kim, et al., Volume regulation and shape bifurcation in the cell nucleus. J. Cell Sci. 128, 3375–85 (2015).

10. J. D. Finan, F. Guilak, The effects of osmotic stress on the structure and function of the cell nucleus. J. Cell. Biochem. 31, 460–467 (2009).

11. Y. Li, et al., Moving Cell Boundaries Drive Nuclear Shaping during Cell Spreading. Biophys. J. 109, 670–686 (2015).

12. X. Cao, et al., A Chemomechanical Model for Nuclear Morphology and Stresses during Cell Transendothelial Migration. Biophys. J. 111, 1541–1552 (2016).

13. F. Alisafaei, D. S. Jokhun, G. V. Shivashankar, V. B. Shenoy, Regulation of nuclear architecture, mechanics, and nucleocytoplasmic shuttling of epigenetic factors by cell geometric constraints. Proc. Natl. Acad. Sci. U. S. A. 116, 13200–13209 (2019).

14. S. B. Khatau, et al., A perinuclear actin cap regulates nuclear shape. Proc. Natl. Acad. Sci. 106, 19017–19022 (2009).

15. D. E. Ingber, Cellular tensegrity: defining new rules of biological design that govern the cytoskeleton. J. Cell Sci. 104 (Pt 3, 613–27 (1993).

16. W. W. Feng, W. H. Yang, On the Contact Problem of an Inflated Spherical Nonlinear Membrane. J. Appl. Mech. 45, 209–214 (1973).

17. S. Balakrishnan, et al., A Nondimensional Model Reveals Alterations in Nuclear Mechanics upon Hepatitis C Virus Replication. Biophys. J. 116, 1328–1339 (2019).

18. M. Velliste, R. F. Murphy, Automated determination of protein subcellular locations from 3D fluorescence microscope images in Biomedical Imaging, 2002. Proceedings. 2002 IEEE International Symposium On, (2002), pp. 867–870.

19. A. Buxboim, et al., Matrix elasticity regulates lamin-A,C phosphorylation and turnover with feedback to actomyosin. Curr. Biol. 24, 1909–1917 (2014).

20. B. A. Danowski, Fibroblast contractility and actin organization are stimulated by microtubule inhibitors. J. Cell Sci. 93, 255–266 (1989).

21. A. Rape, W. H. Guo, Y. L. Wang, Microtubule depolymerization induces traction force increase through two distinct pathways. J. Cell Sci. 124, 4233–4240 (2011).

22. M. S. Kolodney, E. L. Elson, Contraction due to microtubule disruption is associated with increased phosphorylation of myosin regulatory light chain. Proc. Natl. Acad. Sci. U. S. A. 92, 10252–10256 (1995).

23. A. D. Verin, et al., Microtubule disassembly increases endothelial cell barrier dysfunction: Role of MLC phosphorylation. Am. J. Physiol. - Lung Cell. Mol. Physiol. 281, 565–574 (2001).

24. A. A. Birukova, et al., Microtubule disassembly induces cytoskeletal remodeling and lung vascular barrier dysfunction: Role of Rho-dependent mechanisms. J. Cell. Physiol. 201, 55–70 (2004).

25. Y.-C. Chang, P. Nalbant, J. Birkenfeld, Z.-F. Chang, G. M. Bokoch, GEF-H1 Couples Nocodazole-induced Microtubule Disassembly to Cell Contractility via RhoA. Mol. Biol. Cell 19, 2147–2153 (2008).

26. Q. Li, A. Kumar, E. Makhija, G. V. Shivashankar, The regulation of dynamic mechanical coupling between actin cytoskeleton and nucleus by matrix geometry. Biomaterials 35, 961–969 (2014).

27. K. Damodaran, et al., Compressive force induces reversible chromatin condensation and cell geometry–dependent transcriptional response. Mol. Biol. Cell 29, 3039–3051 (2018).

28. D. B. Lovett, N. Shekhar, J. A. Nickerson, K. J. Roux, T. P. Lele, Modulation of nuclear shape by substrate rigidity. Cell. Mol. Bioeng. 6, 230–238 (2013).

29. Y. Merkher, M. B. Alvarez-Elizondo, D. Weihs, Taxol reduces synergistic, mechanobiological invasiveness of metastatic cells. Converg. Sci. Phys. Oncol. 3, 044002 (2017).

30. L. S. Prahl, et al., Microtubule-Based Control of Motor-Clutch System Mechanics in Glioma Cell Migration. Cell Rep. 25, 2591–2604.e8 (2018).

31. D. E. Ingber, Cellular tensegrity : defining new rules of biological design that govern the cytoskeleton. 627, 613–627 (1993).

32. S. Vogel, Comparative biomechanics: life’s physical world (Princeton University Press, 2013).

33. P. Roca-Cusachs, V. Conte, X. Trepat, Quantifying forces in cell biology. Nat. Cell Biol. 19, 742–751 (2017).

34. M. Dembo, Y. L. Wang, Stresses at the cell-to-substrate interface during locomotion of fibroblasts. Biophys. J. 76, 2307–2316 (1999).

35. J. L. Tan, et al., Cells lying on a bed of microneedles: An approach to isolate mechanical force. Proc. Natl. Acad. Sci. 100, 1484–1489 (2003).

36. P. Ngo, P. Ramalingam, J. A. Phillips, G. T. Furuta, Collagen gel contraction assay. Methods Mol. Biol. 341, 103–109 (2006).

37. A. J. Ammit, C. L. Armour, J. L. Black, Smooth-muscle myosin light-chain kinase content is increased in human sensitized airways. Am. J. Respir. Crit. Care Med. 161, 257–263 (2000).

38. M. Uehata, et al., Calcium sensitization of smooth muscle mediated by a Rho-associated protein kinase in hypertension. Nature 389, 990–994 (1997).

39. C. M. Kraning-Rush, J. P. Califano, C. A. Reinhart-King, Cellular traction stresses increase with increasing metastatic potential. PLoS One 7(2012).

40. Y. Margaron, et al., Biophysical properties of intermediate states of EMT outperform both epithelial and mesenchymal states. bioRxiv, 797654 (2019).

41. C. Di Ciano, et al., Osmotic stress-induced remodeling of the cortical cytoskeleton. Am. J. Physiol. - Cell Physiol. 283, 850–865 (2002).

42. T. P. Lele, R. B. Dickinson, G. G. Gundersen, Mechanical principles of nuclear shaping and positioning. J. Cell Biol. 217, 3330–3342 (2018).

43. V. J. Tocco, et al., The nucleus is irreversibly shaped by motion of cell boundaries in cancer and non-cancer cells. J. Cell. Physiol. 233, 1446–1454 (2018).

44. J. Fidorra, T. Mielke, J. Booz, L. E. Feinendegen, Cellular and nuclear volume of human cells during the cell cycle. Radiat. Environ. Biophys. 19, 205–214 (1981).

45. B. Vianay, et al., Variation in traction forces during cell cycle progression. Biol. Cell 110, 91–96 (2018).

46. J. R. Tse, A. J. Engler, Preparation of Hydrogel Substrates with Tunable Mechanical Properties. Curr. Protoc. Cell Biol. 47, 10.16.1-10.16.16 (2010).

47. A. C. Rowat, J. Lammerding, J. H. Ipsen, Mechanical properties of the cell nucleus and the effect of emerin deficiency. Biophys. J. 91, 4649–4664 (2006).

