## Supplementary Information for "Predicting the tension in actin cytoskeleton from the nucleus shape"

Sreenath Balakrishnan[1]

Shilpa R Raju[2]

Anwesha Barua[3]

G.K. Ananthasuresh[2,3]

[1]School of Mechanical Sciences, Indian Institute of Technology, Goa

[2]Department of Mechanical Engineering, Indian Institute of Science, Bengaluru

[3]BioSystems Science and Engineering, Indian Institute of Science, Bengaluru

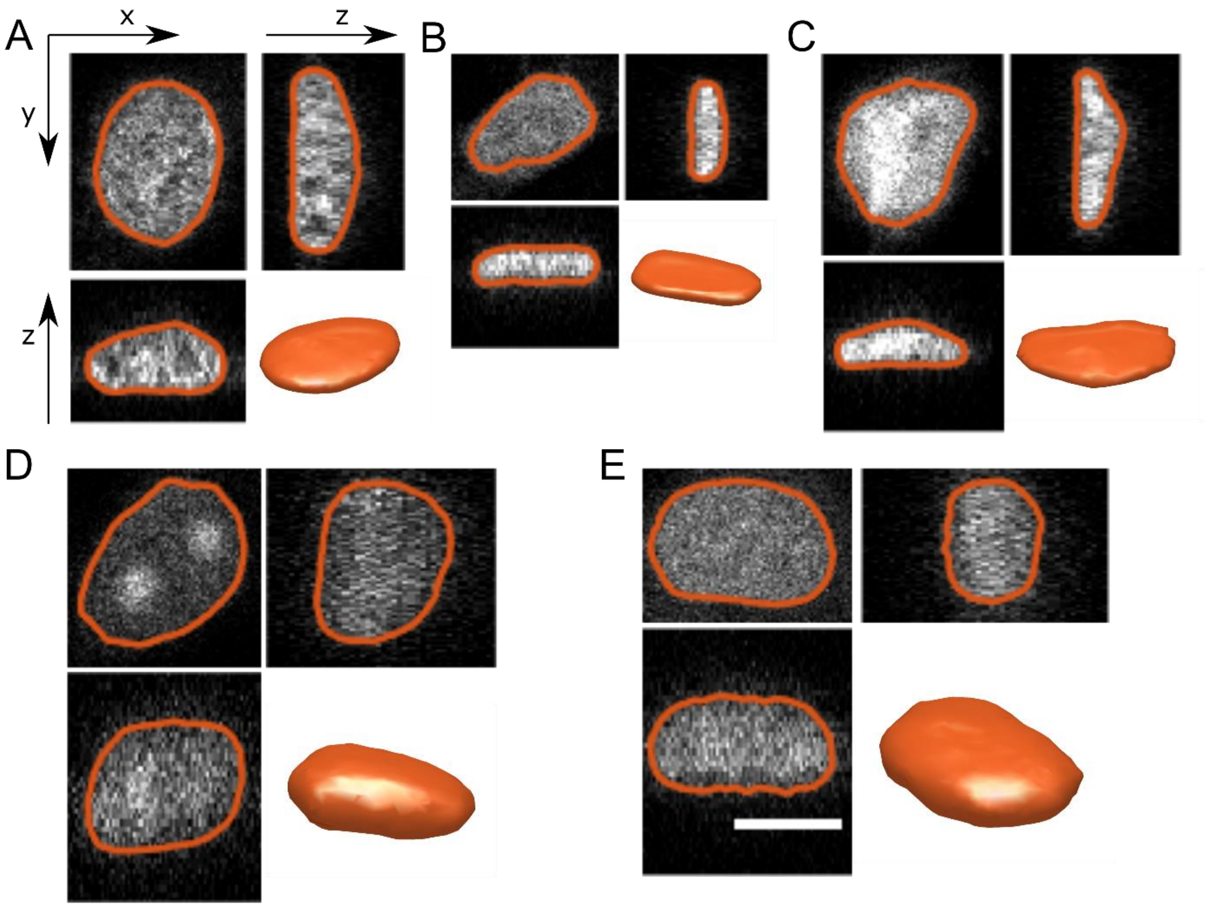


**Figure S1: Obtaining nuclear surface from confocal images.** Three orthogonal sections from the confocal stack of nuclei of (A) 3T3, (B) MDAMB, (C) HeLa, (D) MCF7 and (E) Huh7 cells. The contours of the nuclear surface at these sections and a 3D rendering of the nuclear surface is also shown. Scale bar represents 10 μm.

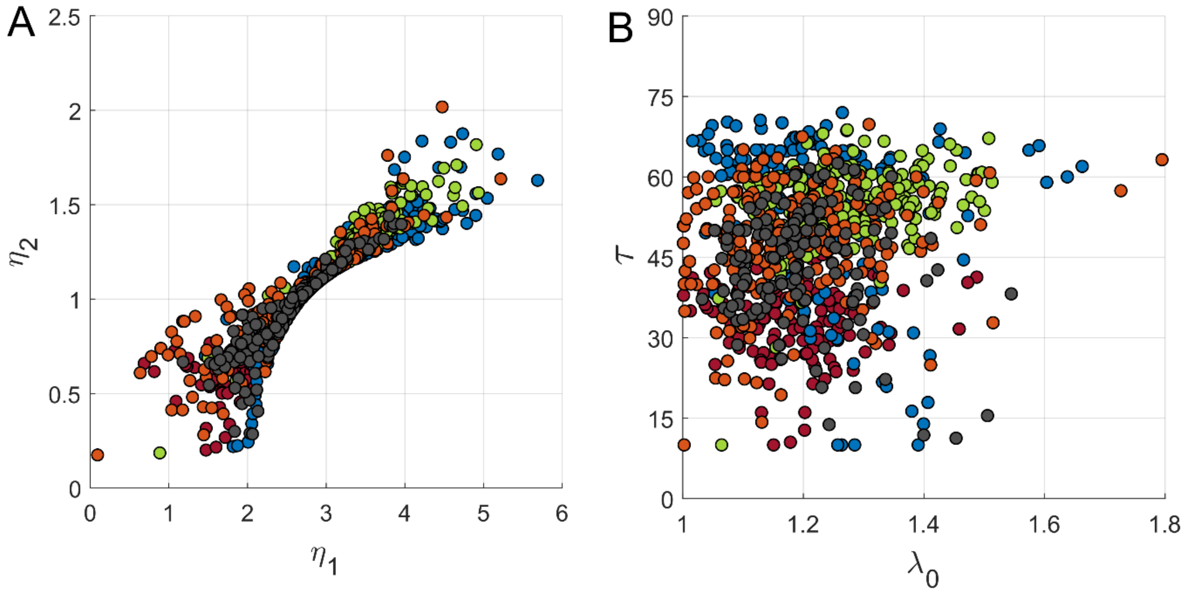


**Figure S2: Correlation between nondimensional parameters**. (A) Scatter plot of $\eta_1$ and $\eta_2$ for MDAMB231 (orange), HeLa (blue), MCF7 (grey), Huh7 (red) and 3T3 (green) cells (B) Scatter plot of $\lambda_0$ and $\tau$.

**Table S1: Correlation between $\lambda_0$, $\tau$ and the components of the nuclear shape parameters along the principal directions, $v_1$ and $v_2$**

| Cell line | $p_{\lambda 0,v1}$ | $p_{\lambda 0,v2}$ | $p_{\tau,v1}$ | $p_{\tau,v2}$ |
|---|---|---|---|---|
| Huh7 | 0.98 | 0.18 | 0.16 | 0.89 |
| HeLa | 0.99 | 0.01 | -0.10 | 0.96 |
| NIH3T3 | 0.99 | -0.03 | 0.47 | 0.84 |
| MDAMB231 | 0.99 | -0.03 | 0.37 | 0.87 |
| MCF7 | 0.96 | -0.22 | 0.07 | 0.95 |

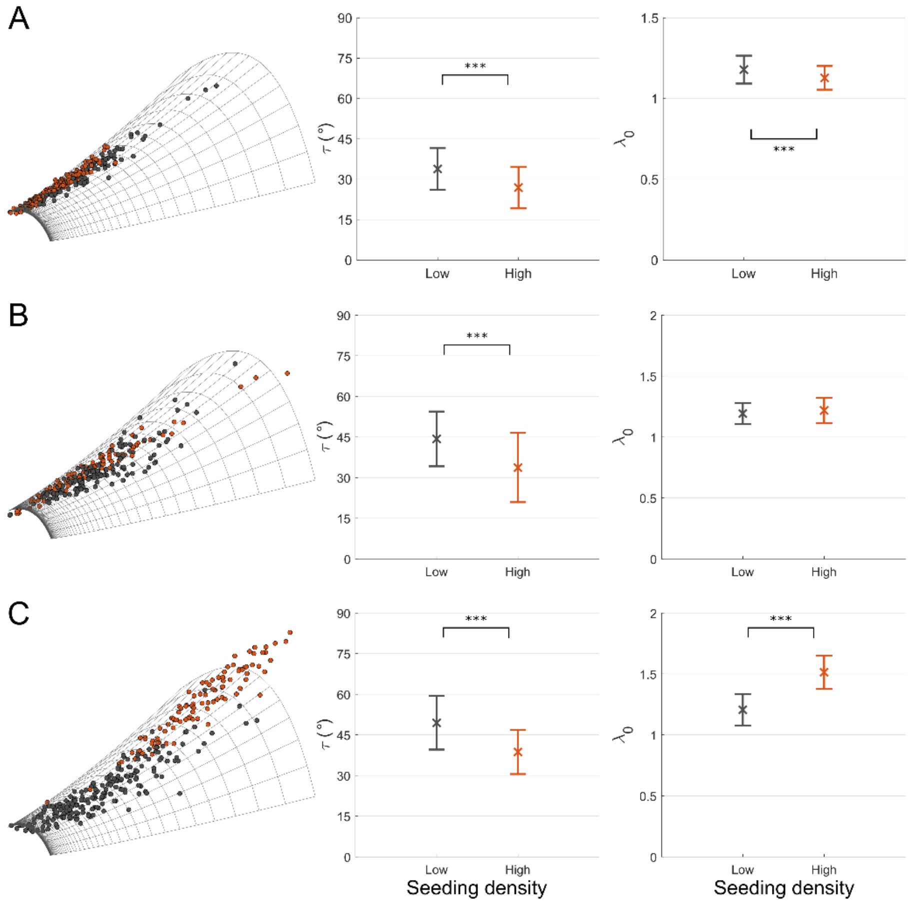


**Figure S3: Comparison of nondimensional parameters at low and high cell-seeding density**. Huh7 (A), MCF7 (B), and MDAMB231 (C) cells were cultured at low and high seeding densitites. Individual nuclei from these cultures were fit our model, and $\lambda_0$ and $\tau$ were estimated. Scatter plot of nuclear shape parameters on the model surface is shown in the left column. Each dot is an individual nucleus and the colors represent black – low density, and red – high density cultures. Bar graphs with the mean and standard deviation of $\tau$ and $\lambda_0$ are shown in the center and right columns respectively. Statistical analysis was performed using Kolmogrov-Smirnov test and significance levels are represented by * $p<0.05$, ** $p<0.01$, and *** $p<0.001$

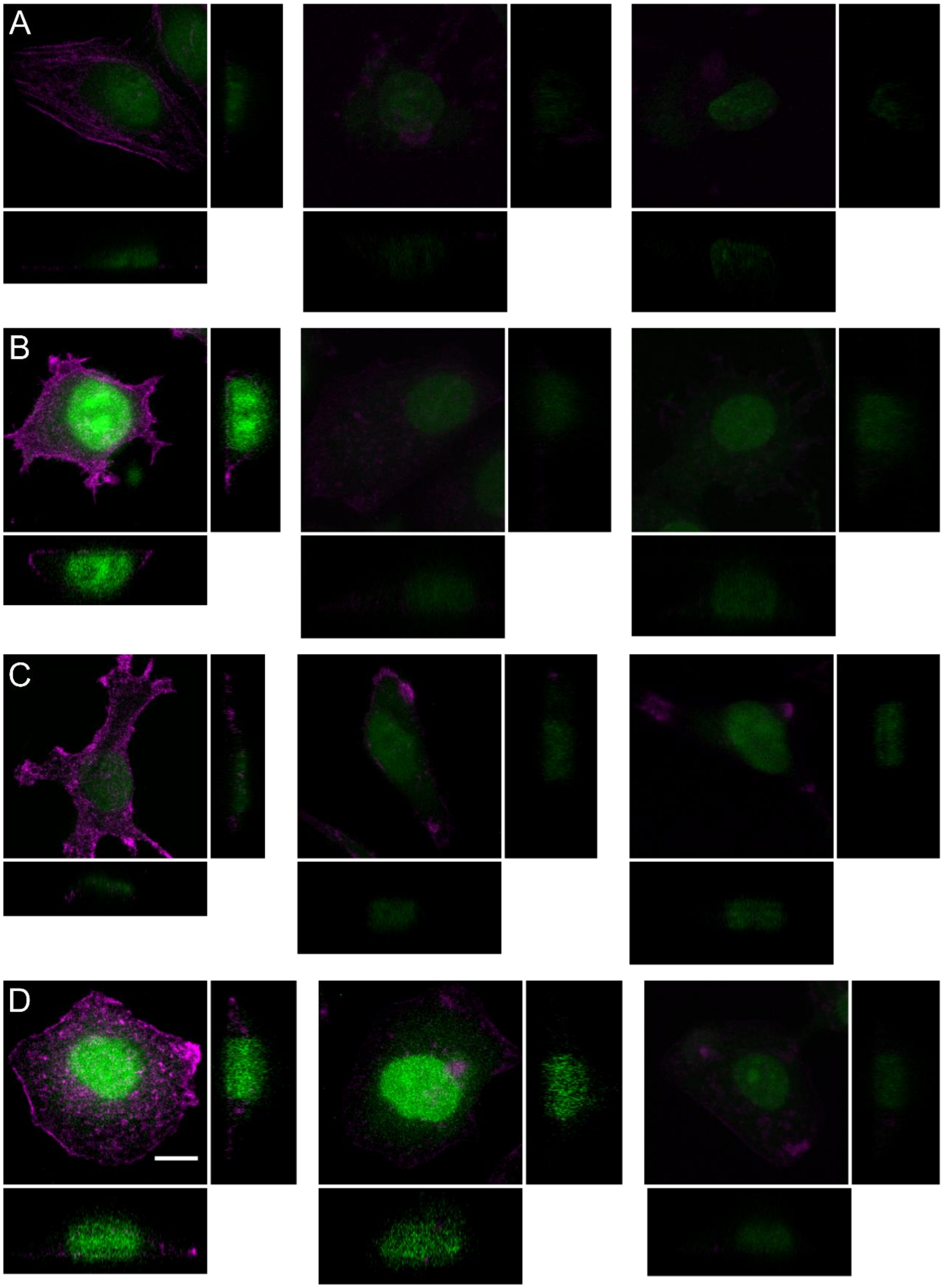


**Figure S4: Confocal stacks of cells treated with Cytochalasin D**. Orthogonal projections of (A) HeLa, (B) MCF7, (C) MDAMB231, and (D) Huh7 cells treated with Cytochalasin D with nucleus - green and actin - magenta. From left to right the columns are control, 0.46 μM, and 0.92 μM Cytochalasin D. The scale bar represents 10 μm

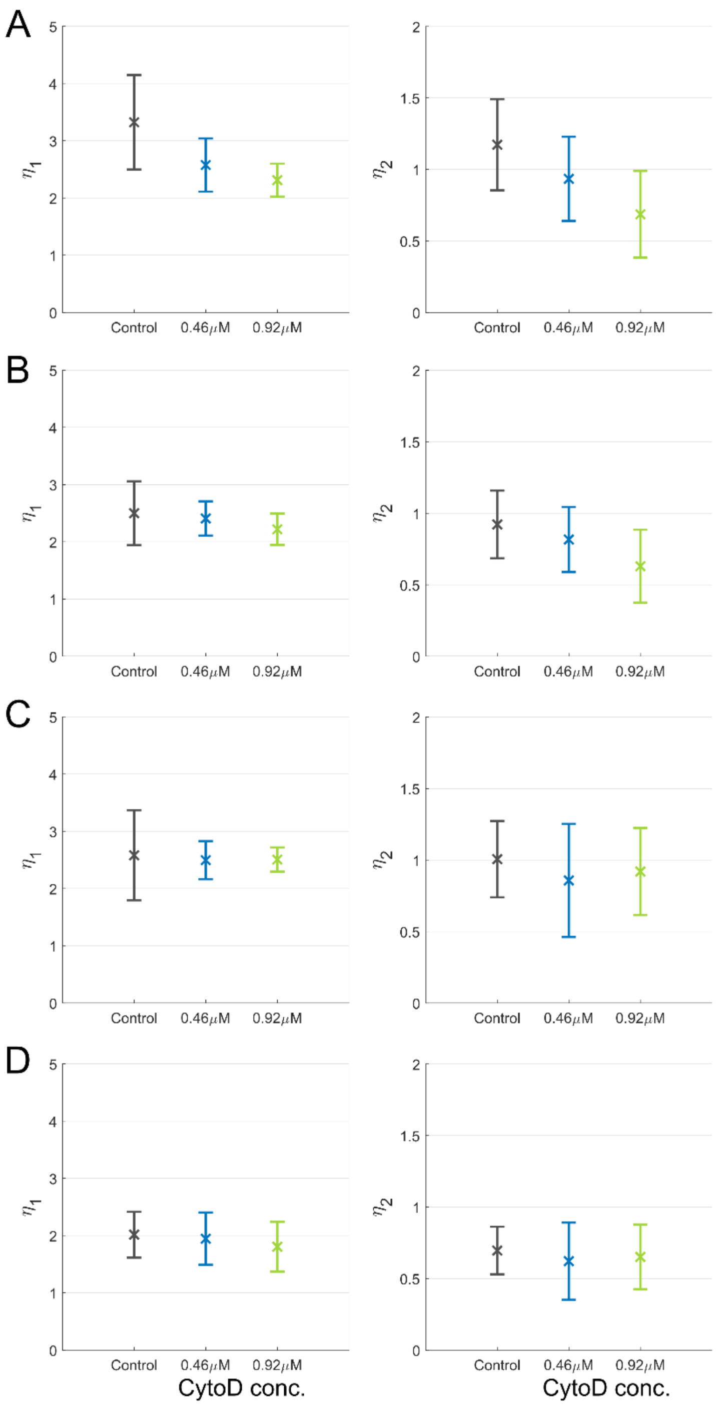


**Figure S5: Changes in $\eta_1$ and $\eta_2$ due to Cytochalasin D treatment**. Four cell lines, HeLa (A), MCF7 (B), MDAMB231 (C), and Huh7 (D) were treated with twp concentrations, 0.46 μM and 0.92 μM, of Cytochalasin D. By fitting our model to individual nuclei, the nondimensional parameters were estimated. Bar graphs with the mean and standard deviation of $\eta_1$ and $\eta_2$ are shown in the left and right columns respectively. The colors represent black – control, blue - 0.46 μM, and green - 0.92 μM of Cytochalasin D.

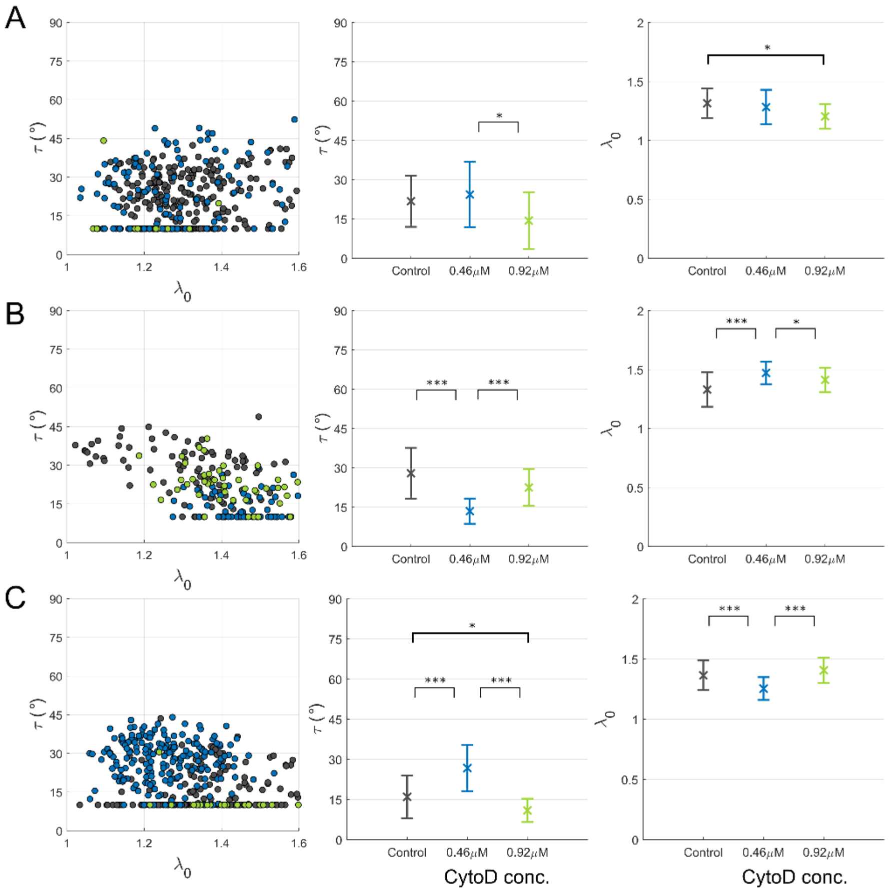


**Figure S6: Changes in nondimensional parameters due to Cytochalasin D treatment on cells with low actin cytoskeletal tension**. (A) MDAMB231, (B) MCF7, and (C) 3T3 cells were cultured at high seeding density to obtain cells with low actin cytoskeletal tension. These cells were treated with two concentrations, 0.46 μM, and 0.92 μM, of Cytochalasin D. By fitting our model to individual nuclei, nondimensional parameters $\lambda_0$ and $\tau$ were estimated. Scatter plot of $\lambda_0$ and $\tau$ is shown in the left column. Each dot is an individual nucleus and the colors represent black – control, blue - 0.46 μM, and green - 0.92 μM of Cytochalasin D. Bar graphs with the mean and standard deviation of $\tau$ and $\lambda_0$ are shown in the center and right columns respectively. Statistical analysis was performed using ANOVA with Bonferroni correction and significance levels are represented by * $p<0.05$, ** $p<0.01$, and *** $p<0.001$

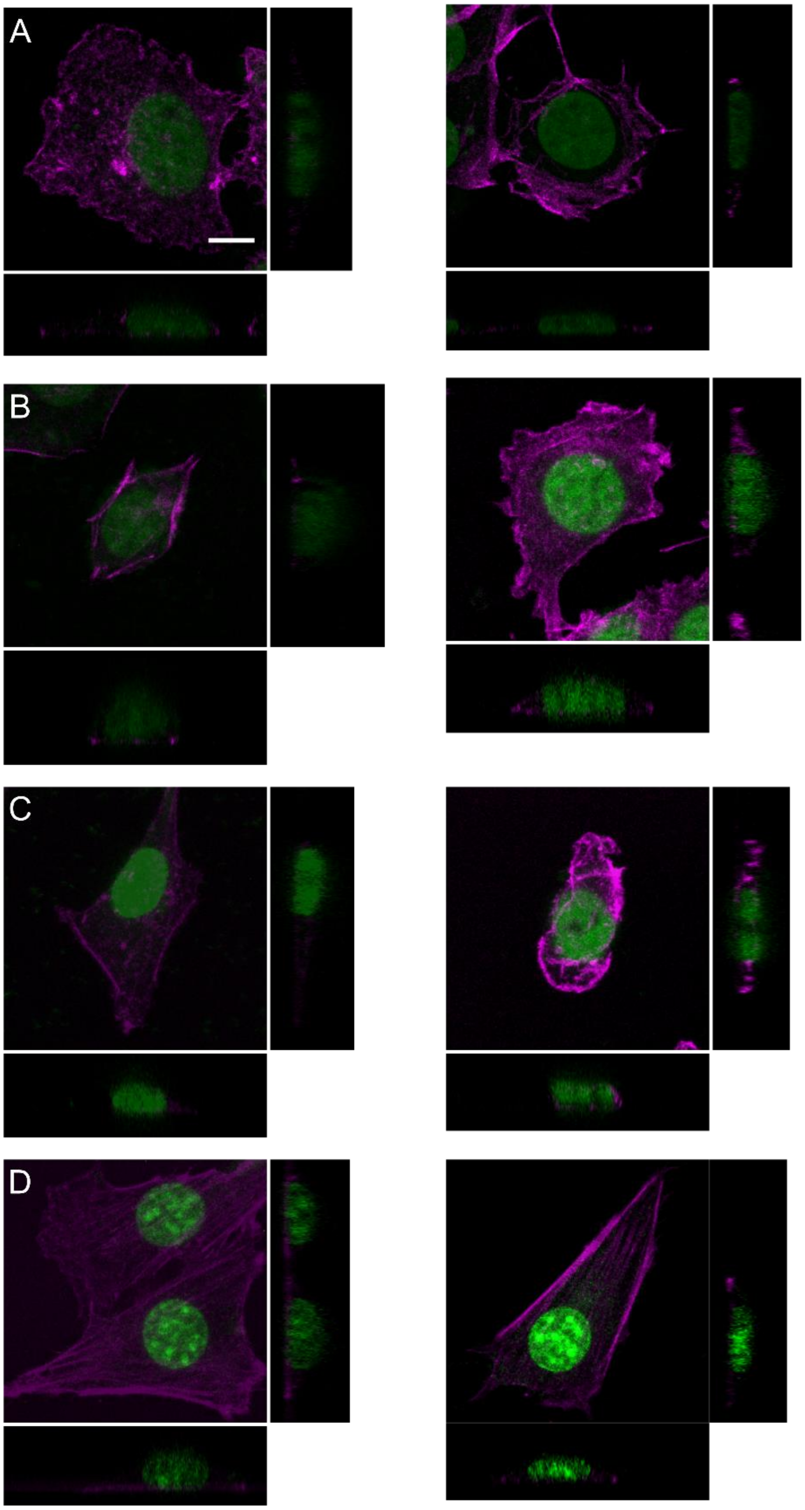


**Figure S7: Confocal stacks of cells treated with Nocodazole**. Orthogonal projections of (A) Huh7, (B) MCF7, (C) MDAMB231, and (D) 3T3 cells treated with Nocodazole with nucleus - green and actin - magenta. The left column is control and the right column is treated with 6 µM Nocodazole. The scale bar represents 10 µm

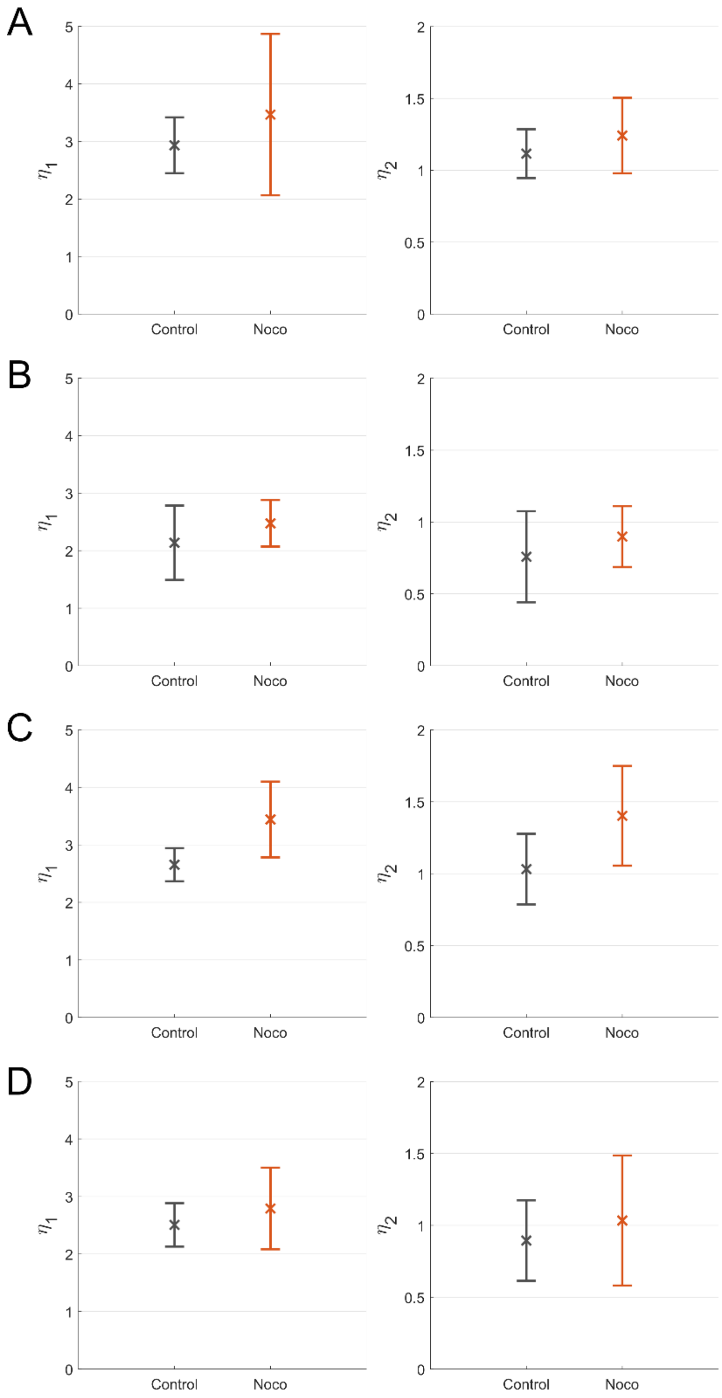


**Figure S8: Changes in $\eta_1$ and $\eta_2$ due to Nocodazole treatment**. Four cell lines, (A) Huh7, (B) MCF7, (C) MDAMB231, and (D) 3T3, were treated with 6 μM Nocodazole. By fitting our model to individual nuclei, the nondimensional parameters were estimated. Bar graphs with the mean and standard deviation of $\eta_1$ and $\eta_2$ are shown in the left and right columns respectively. The colors represent black – control, and red – Nocodazole-treated cells. Bar graphs with the mean and standard deviation of $\eta_1$ and $\eta_2$ are shown in the left and right columns respectively

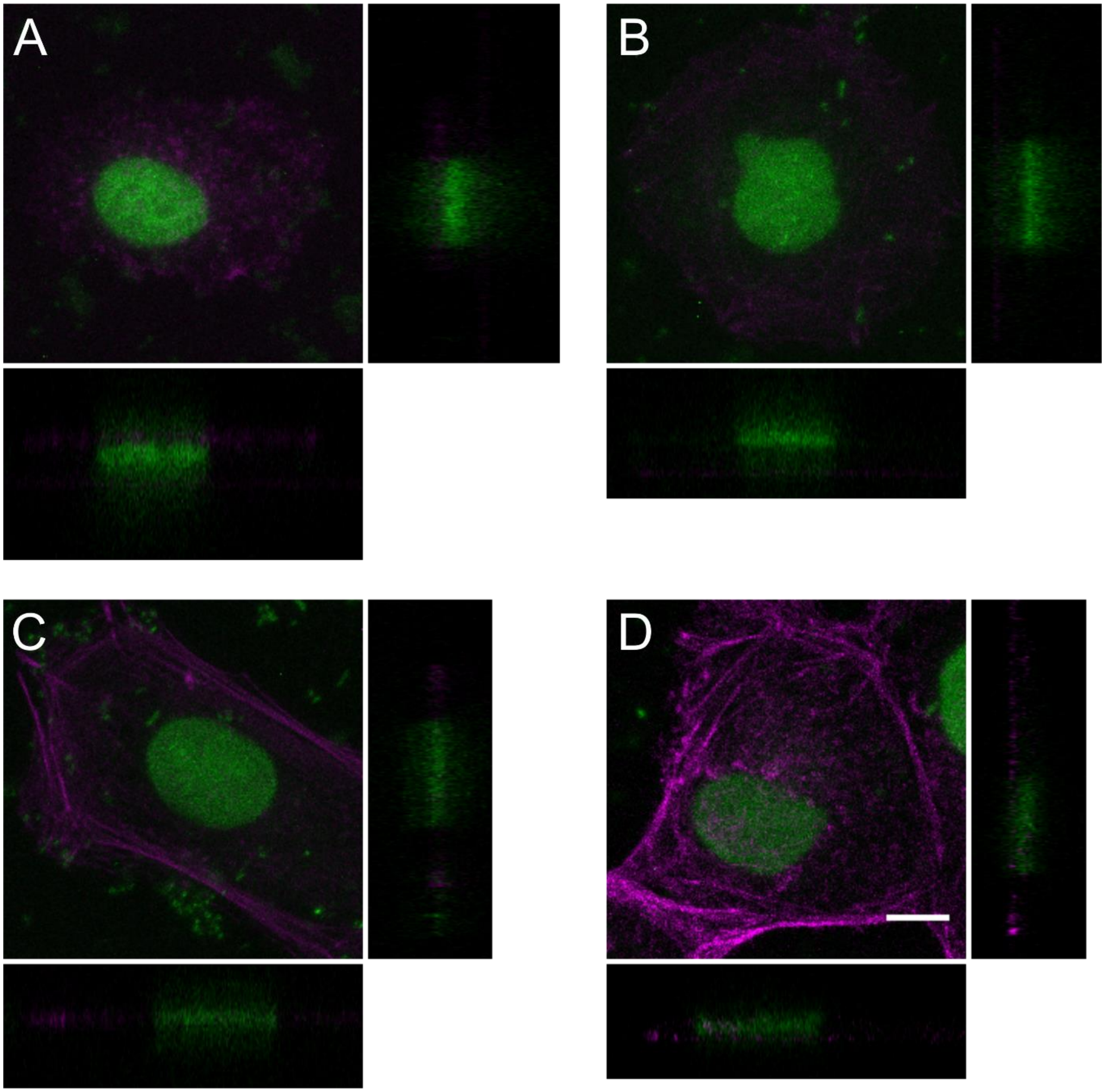


**Figure S9: Confocal stacks of Huh7 cells on substrates of varying elastic modulus**. Orthogonal projections of Huh7 cells on polyacrylamide gels of elastic modulus (A) 2.5 kPa, (B) 11 kPa, and (C) 36 kPa. Huh7 cells on glass coverslip (D) was used as control. The cells were stained for nucleus - green and actin - magenta. The scale bar represents 10 μm

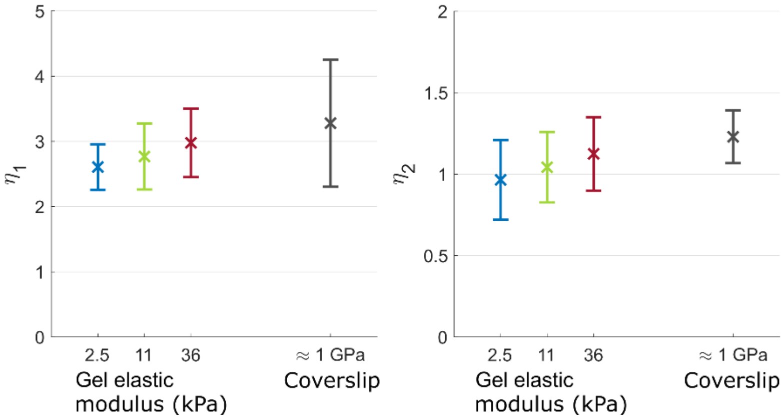


**Figure S10: Changes in $\eta_1$ and $\eta_2$ on varying elastic modulus of substrate**. Huh7 cells were cultured on polyacrylamide gels with elastic modulus of 2.5, 11, and 36 kPa. By fitting our model to individual nuclei, nondimensional parameters $\lambda_0$ and $\tau$ were estimated. Bar graphs with the mean and standard deviation of $\eta_1$ and $\eta_2$ are shown in the left and right columns respectivelyThe colors represent black – control (coverslip ≈ 1 GPa), blue – 2.5 kPa, green – 11 kPa and red – 36 kPa. Bar graphs with the mean and standard deviation of $\eta_1$ and $\eta_2$ are shown in the left and right columns respectively

### Geometric approximation of simulated nuclear shapes from the model

We observed that the simulated nuclear morphologies from our model could be approximated by a simple 'pancake geometry'. The cross-section of this axisymmetric geometry is shown in Fig. S10A. We used this approximation to derive a method for obtaining the aspect ratio, height to diameter, of the nucleus from nuclear volume and projected area. $\tau$ could be further estimated from this aspect ratio. For the simplified geometry, the projected area is

$$A_p = \pi(r+a)^2 \tag{S1}$$

Aspect ratio is

$$\gamma = \frac{a}{r+a} \tag{S2}$$

The projected area can be expressed in terms of the aspect ratio as

$$A_p = \frac{\pi a^2}{\gamma^2} \tag{S3}$$

The expression for the volume is

$$V = \pi a\left(2r^2 + \pi r a + \frac{4}{3}a^2\right) \tag{S4}$$

This relation can be rearranged to facilitate expression in terms of the aspect ratio as

$$V = 2\pi a^3\left[\left(\frac{r}{a}+1\right)^2 - \left(2-\frac{\pi}{2}\right)\left(\frac{r}{a}+1\right) + \left(\frac{5}{3}-\frac{\pi}{2}\right)\right] \tag{S5}$$

Substituting the expression for aspect ratio, Eq. (S2) in Eq. (S5), and dividing by $A_p^{3/2}$,

$$\left(\frac{5}{3}-\frac{\pi}{2}\right)\gamma^3 - \left(2-\frac{\pi}{2}\right)\gamma^2 + \gamma - Q = 0 \tag{S6}$$

where $Q = \left(V\sqrt{\pi}\right)/\left(2A_p^{3/2}\right)$. For small values of $\tau \leq 30°$, the error in estimation of aspect ratio using this expression can be as high as 5% (Fig. S10B). This error was corrected when estimating $\tau$ from aspect ratio. The correction was done by mapping the aspect ratio obtained from this expression to $\tau$ estimated by fitting our model to nuclear volume and projected area (Fig. 6C). Hence, the aspect ratio on the x-axis of Fig. 6C was obtained by solving Eq. (S6) for a given nuclear volume and projected area and $\tau$ on y-axis was estimated by fitting the model.

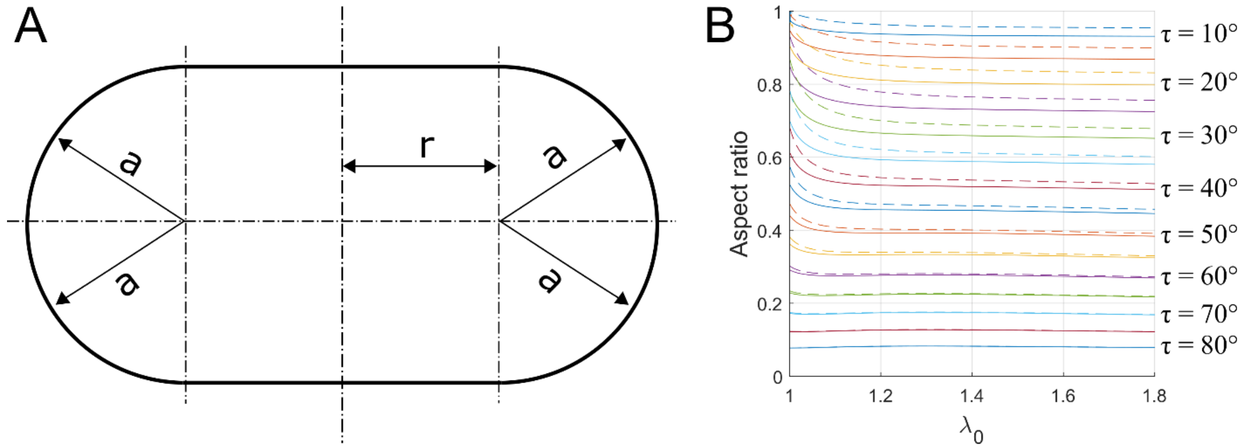


**Figure S11: Approximate method for estimating τ using a simplified geometry**. (A) Nucleus morphology obtained from the model was approximated by an axisymmetric surface with a 'pancake' geometry with circular ends. (B) Comparison of aspect ratio of the nucleus obtained from the model (solid lines) and this approximate geometry (dashed lines).
