## Supplementary material for "Predicting the tension in actin cytoskeleton from the nucleus shape": Obtain_eta1_eta2_l0_tau.m

```
% This code is for calculating the non-dimensional parameters eta_1, eta_2,
% l0 and tau from nuclear shape parameters of individual nuclei
% Sreenath Balakrishnan 2020

%% Header
clearvars; 
close all; 
clc;

%% Parameters
R=5.5; %Radius of the initial sphere for normalizing

%% Data from model simulations
v = xlsread('ModelData.xlsx','v');   % Normalised Volume
a_s = xlsread('ModelData.xlsx','a_s'); % Normalised Surface area
a_p = xlsread('ModelData.xlsx','a_p'); % Normalised Projected area
l0  = xlsread('ModelData.xlsx','l_0'); % l0 = lambda0
tau = xlsread('ModelData.xlsx','tau'); % tau in degrees
eta_1 = xlsread('ModelData.xlsx','eta_1'); % eta1 = (P*R)/(2*E_1*H)
eta_2 = xlsread('ModelData.xlsx','eta_2'); % eta2 = sqrt(F/(P*pi*R^2))

%% load your morphology data here instead of the sample data
% Sample morphology data
A_p = xlsread('MorphologySampleData.xlsx','B2:B11'); %Projected area
A_s = xlsread('MorphologySampleData.xlsx','C2:C11'); %Projected area
V = xlsread('MorphologySampleData.xlsx','D2:D11'); %Volume

%% Initialise variables
n_data = length(V);

% nondimensional parameters for the respective nuclear morphology
eta_1d = zeros(n_data,1); eta_2d = zeros(n_data,1);
l0d = zeros(n_data,1);    taud = zeros(n_data,1);
% error in fitting
er = zeros(n_data,3);

%% Code
% normalize the nuclear shape parameters
vd = V/R^3; a_pd = A_p/R^2; a_sd = A_s/R^2;

for ndata = 1:length(vd)
    % calculating the relative error
    da_p = (a_p - a_pd(ndata))./a_pd(ndata); 
    dv = (v - vd(ndata))./vd(ndata); 
    da_s = (a_s - a_sd(ndata))./a_sd(ndata); 

    % finding the index with least error
    [~,indx]= min(da_p(:).^2 + dv(:).^2 + da_s(:).^2);
    [rw,cl]=ind2sub(size(a_p),indx);
    
    % saving the error and corresponding nondimensional parameters
    er(ndata,:) = [da_p(rw,cl) dv(rw,cl) da_s(rw,cl)];
    eta_1d(ndata) = eta_1(rw,cl); eta_2d(ndata) = eta_2(rw,cl);
    l0d(ndata) = l0(rw,cl); taud(ndata) = tau(rw,cl);
end
```
