## Supplementary material for "Predicting the tension in actin cytoskeleton from the nucleus shape": plot_nuclei_over_model_surface.m

```
% This code is for plotting the surface predicted by the two-parameter
% model and nuclear shape parameters of individual nuclei over it
% Sreenath Balakrishnan 2020

%% Header
clearvars; 
close all; 
clc;

%% Parameters
R=5.5; %Radius of the undeformed sphere for normalizing

%% Data from model simulations
v = xlsread('ModelData.xlsx','v');   % Normalised Volume
a_s = xlsread('ModelData.xlsx','a_s'); % Normalised Surface area
a_p = xlsread('ModelData.xlsx','a_p'); % Normalised Projected area

%% load your morphology data here instead of the sample data
% Sample morphology data
A_p = xlsread('MorphologySampleData.xlsx','B2:B11'); %Projected area
A_s = xlsread('MorphologySampleData.xlsx','C2:C11'); %Projected area
V = xlsread('MorphologySampleData.xlsx','D2:D11'); %Volume

%% Code
% normalize the nuclear shape parameters
vd = V/R^3; a_pd = A_p/R^2; a_sd = A_s/R^2;

% plot the surface predicted by the model
figure; hold on;
surf(a_p,a_s,v,'facecolor','none','edgecolor',[0.31,0.31,0.31],'linewidth',0.8);

% plot the morphology data
scatter3(a_pd,a_sd,vd,'markerfacecolor','r','markeredgecolor','k');

% labeling the axes
xlabel('Norm. Projected area'); ylabel('Norm. Surface area');
zlabel('Norm. Volume');
axis equal; grid on;
view(38.75,0);
```
