## Supplementary material for "Predicting the tension in actin cytoskeleton from the nucleus shape": readme.txt

Use the 'plot_nuclei_over_model_surface.m' script to plot the nuclear shape parameters over the model surface. For this, copy your nuclear shape data to 'MorphologySampleData.xlsx', choose a value for R and run the script. Tune R such that more than 95% of the nuclei have lambda0>1.

Now use this tuned R in the 'Obtain_eta1_eta2_l0_tau.m' script to approximately fit the nondimensional parameters. 
